# Modified h*CFTR* mRNA restores normal lung function in a mouse model of cystic fibrosis

**DOI:** 10.1101/202853

**Authors:** AKM Ashiqul Haque, Alexander Dewerth, Justin S. Antony, Joachim Riethmüller, Ngadhnjim Latifi, Hanzey Yasar, Petra Weinmann, Nicoletta Pedemonte, Elvira Sondo, Julie Laval, Patrick Schlegel, Christian Seitz, Brigitta Loretz, Claus-Michael Lehr, Rupert Handgretinger, Michael S. D. Kormann

## Abstract

Being a classic monogenic disease, gene therapy has always been a promising therapeutic approach for Cystic Fibrosis (CF). However, numerous trials using DNA or viral vectors encoding the correct protein resulted in a general low efficacy. In the last years, chemically modified messenger RNA (cmRNA) has been proven to be a highly potent, pulmonary effective drug. We thus explored the expression of human (h)CFTR encoded by h*CFTR* cmRNA *in vitro*, analyzed by flow cytometry and Western Blot and its function with a YFP assay. Very similar effects could be observed *in vivo* when h*CFTR* cmRNA was assembled with Chitosan-coated PLGA to nanoparticles (NPs) and intratracheally (i.t.) or intravenously (i.v) injected, the latter one as an alternative administration route to circumvent the clogged airways of CF patients. This significantly improved lung function, which suggests that h*CFTR* cmRNA-NPs are a promising therapeutic option for CF patients independent of their *CFTR* genotype.

## Introduction

Cystic fibrosis (CF), the most common life-limiting autosomal-recessive disease in Caucasian populations (1/2,500 newborns), affects more than 80,000 people world-wide (1). It is caused by different mutations within the gene encoding for the CF transmembrane and conductance regulator (CFTR). Those mutations result in impaired anion secretion and hyper-absorption of sodium ions across epithelia (2, 3). Chronic lung disease and slow lung degradation is the major factor contributing to both the mortality and a strongly reduced quality of life (4, 5). With currently available therapies, the mean survival is between 35 and 45 years (6, 7). Since the CFTR gene was first cloned in 1989, many efforts have been made to deal with the mutations at a cellular and genetic level (8). Gene therapy approaches made it quickly to the clinic aiming to deliver viral CFTR encoding vectors [such as adenoviruses (Ad) or adeno-associated viruses (AAV)] to CF patients (9, 10). However, none of the clinical studies and current treatments seem to provide sufficient human (h)CFTR expression to prevent the ultimately lethal CF symptoms in the respiratory tract of CF patients. Furthermore, repeated administration of viral vectors or DNA lead to the development of unwanted immune reactions, mainly due to viral capsids and vector-encoded proteins (11–13). Newly designed viral vectors circumvent those problems and can be administered repeatedly, but from a clinical perspective the field is still in need of a therapeutic tool that combines efficient expression in lungs and other (affected) organs and cells, while avoiding immunogenicity and genotoxicity completely (14–16). Recently, *in vitro* transcribed (IVT) chemically modified messenger RNA (cmRNA) came into focus, which has the potential to combine the mentioned advantages in a single-stranded molecule (17–19). Chemically modified mRNA has been tested for repeated administration, without developing immune responses or losing efficacy, presenting h*CFTR* cmRNA complexed with biodegradable chitosan-coated PLGA nanoparticles (NPs) as a promising therapeutic for the treatment of CF patients (1, 18, 20). Versatile delivery options of mRNA ensures unique possibility to utilize it in early infants as well as in adults, independent of the underlying *CFTR* mutation. To best of our knowledge, we provide the first *in vivo* studies delivering h*CFTR* cmRNA to the lungs of CFTR deficient mice by intravenous (i.v.) and intratracheal (i.t.) administration, complexed with NPs. We hereby demonstrate a proof of concept of cmRNA-NP-mediated, ELISA quantified hCFTR expression in the lungs of *Cftr*^-/-^ mice, leading to significantly reduced chloride secretion and, more importantly, restored normal lung function parameters.

## Materials and Methods

### mRNA production

h*CFTR* was PCR amplified from pcDNA3.hCFTR with the fusion of *Kpn*l and *Eco*RI restriction sites and cloned into a polyA-120 containing pVAX (pVAX.A120, www.lifetechnologies.com) by sticky-end ligation using the mentioned restriction sites. For control experiments, DsRED reporter protein was sub-cloned into pVAX.A120 vector from its original vector pDsRED (www.clontech.com). For *in vitro* transcription (IVT), the plasmids were linearized downstream of the poly-A tail with *Xho*l (www.neb.com). IVT reaction was carried out using MEGAscript T7 Transcription kit (www.ambion.com) with an anti-reverse CAP analog (ARCA) at the 5’ end (www.trilink.com). To produce modified mRNA, the following chemically modified nucleosides were added to the IVT reaction in the indicated ratios: uridine-tri- phosphate (UTP) and cytidine-tri-phosphate (CTP) were fully replaced by N1-Pseudo-h *CFTR* UTP and 5-Methyl-CTP, abbreviated to 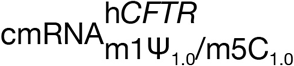 and partly replaced by the incorporation of 25% 2-Thio-UTP and 25% 5-methyl-CTP, respectively, abbreviated to 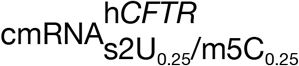 (www.trilink.com). The h*CFTR* and *DsRed* mRNA were purified using the MEGAclear kit (www.ambion.com) and analyzed for size and concentration using a RNA NanoChip 6000 for Agilent 2100 Bioanalyzer (www.agilent.com).

### Mammalian cell culture and transfection

Human bronchial epithelial (HBE) and cystic fibrosis epithelial (CFBE) cell lines were maintained in Minimum Essential Medium (MEM, www.biochrom.com) supplemented with 10% (v/v) heat-inactivated Fetal Calf Serum, L-Glutamine (2 mM) and Penicillin-Streptomycin (50 U/ml). Cells were incubated at 37 °C in a humidified atmosphere containing 5% CO_2_ until they reached 80-90% confluency. Cell lines were washed with cold sterile PBS and detached by trypsin-EDTA. Trypsinisation was stopped by adding MEM medium with serum. Cells were collected and spun down at 500 × g for 5 minutes before resuspension in fresh MEM. One day before transfection 250,000 cells/well/1 ml were plated in 12-well plates and grown overnight in MEM without antibiotics. At a confluency of 70-90%, cells were then transfected with 1000 ng mRNA encoding hCFTR using lipofectamine 2000 (www.invitrogen.com) following the manufacturer’s instructions and after changing the media to the reduced serum media, Opti-MEM (www.thermofisher.com). After 5 h, cells were washed with PBS and serum-containing MEM was added. Cells were kept for 24 h and 72 h before further analyses.

### Flow cytometry analyses

All flow cytometry analyses were performed using a Fortessa X-20 (www.bdbioscience.com). For detection of hCFTR protein in HBE and CFBE cell lines, cells were transfected as described above and subsequently prepared for intracellular staining using a Fixation/Permeabilization Solution Kit as directed in the manufacturer’s instruction (www.bdbioscience.com). As primary antibody mouse anti-human hCFTR clone 596 (1:500, kindly provided by the cystic fibrosis foundation therapeutics Inc.) has been used. As secondary antibody served Alexa Fluor 488 goat anti-mouse IgG (1:1,000, www.lifetechnologies.com). At least 20,000 gated cells per tube were counted. Data were analyzed with FlowJo software, version 10.

### YFP-based functional assay

CFTR activity following transient transfection of h*CFTR* (c)mRNA in A549 cells was determined using the halide-sensitive yellow fluorescent protein YFP-H148Q/I152L (21). CFTR deficient A549 cells stably expressing the YFP were plated in 96-well microplates (50,000 cells/well) in 100 μl of antibiotic-free culture medium and, after 6 h, transfected with either plasmids carrying the coding sequence for CFTR or different h*CFTR* (c)mRNA. For each well, 0.25 μg of mRNA or plasmid DNA and 0.25 μl of Lipofectamine 2000 were pre-mixed in 10 μl of OPTI-MEM (www.invitrogen.com) to generate transfection complexes that were then added to the cells. After 24 hours, the complexes were removed by replacement with fresh culture medium. The CFTR functional assay was carried out 24, 48, or 72 h after transfection. For this purpose, the cells were washed with PBS and incubated for 20-30 min with 60 μl PBS containing forskolin (20 μM). After incubation, cells were transferred to a microplate reader (FluoStar Galaxy; www.bmg.labtech.com) for CFTR activity determination. The μlate reader was equipped with high-quality excitation (HQ500/20X: 500 ± 10 nm) and emission (HQ535/30M: 535 ± 15 nm) filters for yellow fluorescent protein (www.chroma.com). Each assay consisted of a continuous 14-s fluorescence reading (5 points per second) with 2 s before and 12 s after injection of 165 μl of a modified PBS containing 137 mM NaI instead of NaCl (final NaI concentration in the well: 100 mM). To determine iodide (I^−^) influx rate, the final 11 s of the data for each well were fitted with an exponential function to extrapolate initial slope. After background subtraction, cell fluorescence recordings were normalized for the initial average value measured before addition of I^−^. For each well, the signal decay in the final 11 s of the data caused by YFP fluorescence quenching was fitted with an exponential function to derive the maximal slope that corresponds to initial influx of I^−^ into the cells (21). Maximal slopes were converted to rates of variation of intracellular I^−^ concentration (in mM/s) using the equation: d[I^−^]/dt = K_l_[d(F/F_0_)/dt. Where K_l_ is the affinity constant of YFP for I^−^, and F/F_0_ is the ratio of the cell fluorescence at a given time vs. initial fluorescence (21).

### Whole blood assay

Blood from three different, healthy donors was taken and collected in EDTA collection tubes (www.sarstedt.com). For each treatment group 2 ml of EDTA-blood was transferred into 12-well plates and treated accordingly. R-848 (Resiquimod, www.sigmaaldrich.com) was added at a concentration of 1 mg/ml to the respective blood positive controls. (Un-)modified h*CFTR* mRNA and pDNA (15 μg each) were complexed to NPs at a ratio of 1:10. Samples were incubated at 37 °C in a humidified atmosphere containing 5% CO_2_. At 6h and 24h, 1ml of whole blood was transferred into micro tubes containing serum gel (www.sarstedt.com) and spun down at 10,000 × g for 5 min to obtain serum. Sera were stored at −20 °C for further cytokine measurement analyses.

### Animal experiments

All animal experiments were approved by the local ethics committee and carried out according to the guidelines of the German Law for the Protection of Animals (file number: 35/9185.81-2 / K/16). *Cftr*^-/-^ mice (CFTR^tm1Unc^) were purchased from Jackson Laboratory (www.jax.org) at an age of 6 to 8 weeks and were maintained under standardized specific pathogen-free conditions on a 12 h light-dark cycle. Food, water as well as nesting material were provided *ad libitum*. Prior to i.t. spray applications, mice were anesthetized intraperitoneally (i.p.) with a mixture of medetomidine (0.5 mg/kg), midazolam (5 mg/kg) and fentanyl (50 μg/kg). *Cftr*^-/-^ mice received 20 μg or 40 μg of h*CFTR* (c)mRNA or equivalent of 20 μg or 40 μg (calculated using nmols) h*CFTR* pDNA encapsulated in chitosan-coated PLGA nanoparticles [Chitosan (83% deacetylated (Protasan UP CL 113) coated PLGA (poly-D,L-lactide-co- glycolide 75:25 (Resomer RG 752H) nanoparticles; short: NPs] by intratracheal (i.t.) spraying (n=4), and intravenous (i.v.) injection (n=4−7) into the tail vein. Mock treated control *Cftr*^-/-^ mice received 20 μg *DsRed* mRNA complexed to NPs (n=5) by i.t. delivery or just 200 μl of NPs by both i.v. and i.t. delivery. For both interventions, (c)mRNA-NP and pDNA-NP complexes were administered in a total volume of 200 μl. Mice received two injections on a three day interval (day 0 and day 3). Detailed description of the i.t. procedures are explained in previous published study (22). After 6 days mice were sacrificed for further end point analyses. To assess immune responses to (un-)modified h*CFTR* mRNA and h*CFTR* pDNA, C57/BL6 mice (n=4 per group) were treated as described for *Cftr*^-/-^ mice. As positive controls served mice that received *E. coli* mRNA-NPs (20 μg) intravenously. C57BL/6 mice received one injection of 20 μg mRNA complexed to NPs. After 6 h, 24 h and 72 h mice were sacrificed and blood was collected to obtain serum.

### Pulmonary mechanics

Lung function for each group was evaluated using a FlexiVent^®^ (www.scireq.com). Prior to tracheostomy, mice were anaesthetized intraperitoneally as described above. After anaesthesia, a 0.5 cm incision was performed from the rostral to caudal direction. The flap of skin was retracted, the connective tissue was dissected, and the trachea was exposed. The trachea was then cannulated between the second and third cartilage rings with a blunt-end stub adapter. The mouse was connected to the FlexiVent^®^ system and respiratory mechanics were measured.

### Salivary assay

Prior to tracheostomy, anaesthetized mice were injected with 50 μl of 1 mM acetylcholine (ACh) in the cheek to stimulate production of saliva. The fluid was collected via glass capillaries and a chloride assay was performed using the Chloride (Cl^−^) Assay Kit according to the manufacturer’s protocol (www.sigmaaldrich.com). Briefly, saliva was diluted at a ratio of 1:100 with water in a total volume of 50 μl and subsequently 150 μl chloride reagent was added. After 15 min incubation at room temperature in the dark, absorbance was measured at 620 nm using an Ensight Multimode plate reader (www.perkinelmer.com).

### Western blot analysis

Protein lysate isolated from cell lines was separated on Bolt NuPAGE 4-12% Bis-Tris Plus gels and a Bolt Mini Gel Tank (all from www.lifetechnologies.com). Immunoblotting for hCFTR was performed by standard procedures according to the manufacturer’s instructions using the XCell II Mini-Cell and blot modules (www.lifetechnologies.com). After blocking with Blocking buffer (5% Nonfat Dry milk, www.cellSignaling.com) for 1 h at room temperature, primary antibodies against hCFTR (clone 596, 1:1,000) or mouse anti-GAPDH (1:5,000, www.scbt.com) were incubated overnight; horseradish peroxidase-conjugated secondary antibodies (1:5000, anti-mouse from www.dianova.com) were incubated for 1 h at room temperature. Blots were processed using ECL Prime Western Blot Detection Reagents (www.gelifesciences.com). Semiquantitative analysis was performed using the ImageJ software.

### Real-time RT-PCR

After i.t. or i.v. injection of differently modified h*CFTR* cmRNA the lungs were isolated at day 6 (experimental end point) homogenized and lysed with tubes of the Precellys Ceramic Kit 1.4/2.8 mm at 5,000 rpm for 20 s in a Precellys Evolution Homogenizer for subsequent RNA-isolation (all from www.peqlab.com). Reverse transcription of 50 ng RNA was carried out using an iScript cDNA synthesis kit (www.bio-rad.com). Detection of hCFTR mRNA was performed by SYBR-Green based quantitative Real-time PCR in 20 μl reactions on a ViiA7 (www.lifetechnologies.com). In all involved procedures we strictly followed the MIQE protocols for RealTime experiments (23). Pre- and post-reaction rooms were strictly separated. Reactions were incubated for 10 min at 95 °C, followed by 40 cycles of 15 s at 95°C and 2 min at 50°C (annealing and extension), followed by standard melting curve analysis. The following primer pairs were used: hCFTR fwd TGTACGGCTACAGGGGAA, hCFTR rev GCCGATAGGCAGATTGTA; house-keeping gene 18S rRNA fwd GGGAGCCTGAGAAACGGC, 18S rRNA rev GACTTGCCCTCCAATGGATCC.

### Enzyme-linked immunosorbent assays (ELISAs)

To detect protein levels of hCFTR after i.t. or i.v. injection of differently modified h*CFTR* cmRNA, the lungs were isolated at day 6 (experimental end point). A human CFTR ELISA kit was used (www.elabscience.com). Protein was isolated in 600 μl RIPA-buffer and 5 μl protease inhibitor cocktail using the Precellys Ceramic Kit with a bead size of 1,4/2,8 mm (www.sigmaaldrich.com). Tissue was homogenized in a Precellys Evolution Homogenizer at 6,500 rpm for 10 s for a total of three cycles, each interrupted by a 15 s break (www.peqlab.com). Subsequently, supernatants were kept on ice and additionally homogenized 10 times with a 20G needle and incubated for 20 min (www.bdbioscience.com). Lysates were spun down for 20 min at 13,000 × g and 4°C. Supernatant was collected and stored at −20°C for further use. Prior to hCFTR ELISA detection, protein concentration was measured using the Pierce BCA protein assay kit (www.thermofisher.com). For each sample an equal amount of 15 μg whole protein lysate was used. For cytokine measurement, blood from mice and donors was taken to obtain serum and tested for IFN-α and TNF-α production as directed in the manufacturer’s instructions (www.bdbioscience.com).

### Statistics

All analyses were performed using the Wilcoxon-Mann-Whitney test with Graphpad Prism Version 6 (www.graphpad.com). Most of the data are represented as mean ± SD; box plot data are represented as mean ± minimum to maximum values *P* ≤ 0.05 (two-sided) was considered statistically significant.

## Results

### h*CFTR* (c)mRNA and hCFTR protein quantification *in vitro*

To evaluate the influence of chemical nucleoside modification on h*CFTR* mRNA, we first conducted a set of *in vitro* analyses to characterize the efficacy and functionality of hCFTR protein expression. First, we compared the expression profile of plasmid-encoded hCFTR, unmodified h*CFTR* mRNA and two well-defined nucleoside modifications which have been described to exert state-of-the-art stability/expression *in vitro* or in lung-specific cell contexts *in vivo* (1, 24–26). Flow cytometry analyses 24 h after transfection of human cystic fibrosis bronchial epithelial (CFBE) cells showed hCFTR positive cells ranging from 15.8% after h*CFTR* pDNA transfection to 23.7% after unmodifed h*CFTR* mRNA transfection and up to 33.6% and 49.6% after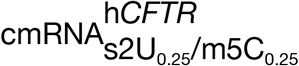 and 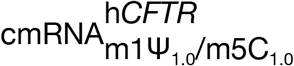ransfecion, resspetivly. At ^24^ h all transfection rates (hCFTR-positive cells, marked as black dots) and hCFTR median fluorescence intensities (MFIs, marked as columns) of unmodifed h*CFTR* mRNA, 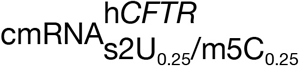, and 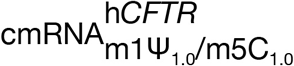, were significantly higher compared to pDNA (*P* ≤ 0.05; Figure 1A, left lower panel).

**Fig. 1:**
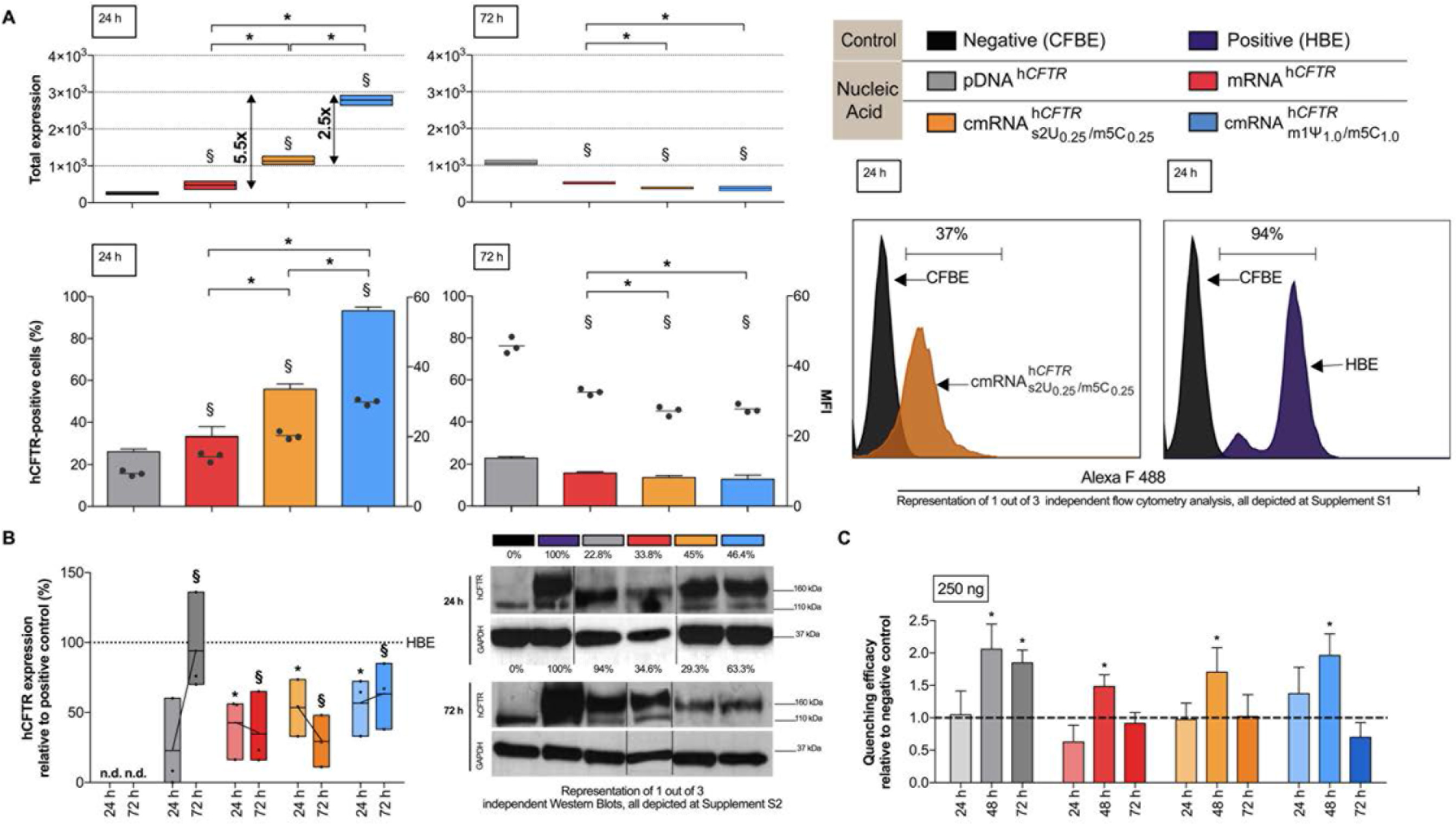
(c)mRNA-mediated expression and function of *hCFTR in vitro*. (**A**) Percentage of hCFTR positive CFBE cells and total expression of hCFTR 24 h and 72 h after transfection with 1 μg h*CFTR* pDNA or (chemically modified) h*CFTR* mRNAs, detected by flow cytometry. ⋆, *P*≤ 0. 05 versus unmodified h*CFTR* mRNA; §, *P* ≤ 0.05 vs. pDNA. (**B**) Western Blots, semi-quantifying human CFTR in the cell cultures used in (**A**), normalized to GAPDH and put relative to CFTR levels in HBE cells. ⋆, *P* ≤ 0.05 versus CFBE controls at 24 h and §, *P* ≤ 0.05 versus CFBE controls at 72 h.(**C**) Quenching efficacy of pDNA or mRNA encoded h*CFTR* in A549 cells relative to untransfected CFBE controls was measured at 24 h, 48 h and 72 h posttransfection. ⋆, *P* ≤ 0.05 versus untransfected controls; MFI, median fluorescence intensities. All other bar graph data are depicted as means ± SDs while box plots data are depicted as the means ± minimum to maximum values.

Total hCFTR expression, defined as median fluorescent intensity (MFI) multiplied by the transfection efficiency, was significantly higher of 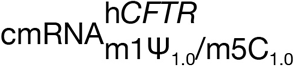, unmodified h*CFTR* mRNA and 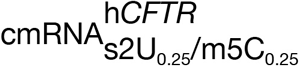 at 24 h compared to pDNA (*P* ≤ 0.05; Figure 1A, left upper panel). In contrast, after 72 h all three h*CFTR* (c)mRNAs expressed significantly lower compared to h*CFTR* pDNA transfected cells, reflected both in percentage of positive cells, MFI and in total hCFTR expression (*P* ≤ 0.05; Figure 1A, h *CFTR* h *CFTR* both right panels). Expression of 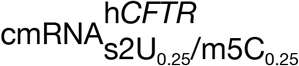 and 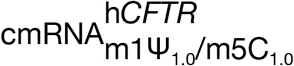 compared to unmodified h*CFTR* mRNA after 72 h was also significantly lower (*P* ≤ 0.05).

To confirm and substantiate those findings, we performed Western blot analyses of protein lysates taken from transfected CFBE cells at 24 h and 72 h post treatment (Figure 1B). As a positive control served protein lysate from untransfected HBE cells, and GAPDH was used to normalize band intensities. At 24 h h*CFTR* pDNA transfected CFBE cells showed an average of 22.8% of the protein expression of hCFTR observed in HBE cells, which increased 4.1-fold to 94.0% at 72 h (Figure 1B). This drastic increase of hCFTR expression after pDNA transfection goes well in line with the observations in flow cytometry as does the quick onset of hCFTR expression after h*CFTR* (c)mRNA transfection at 24 h (Figure 1B). However, relative to the 24 h time- point, hCFTR expression either remained nearly static (unmodified h*CFTR* mRNA resulted in 33.8% and 34.7% expression at 24 h and 72 h, respectively), decreased (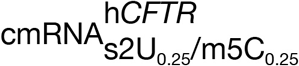 resulted in 45% and dropped to 29.3% hCFTR expression at h*CFTR* 24 h and 72 h, respectively) or increased (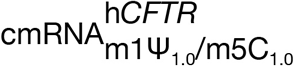, 46.4% at 24 h and raised to 63.3% at 72 h). Ultimately, the expression of h*CFTR* mRNA *in vitro* was strongly dependend on its chemical modification, with 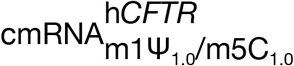 resulting in the most robust hCFTR expression.

### hCFTR (c)mRNA functionality test *in vitro*

For functional analysis of the (c)mRNA-encoded CFTR channel, we performed a YEP-based functional assay using CFTR null A549 cells which stably express halide sensitive YFP-H148Q/I152L (25). Quenching of the YFP signal induced by hCFTR channel-mediated I^−^ influx is reciprocally proportional to hCFTR channel function (21, 27). Figure 1C shows the quenching efficacy after transfection of 250 ng h*CFTR* (c)mRNA, for three different time points, normalized to mock transfected cells. In pDNA transfected cells, the quenching efficacy was significantly higher after 48 h and stayed high even after 72 h (*P* ≤ 0.05), while unmodified as well as modified h*CFTR* mRNA transfected cells revealed a single peak quenching at 48 h (*P* ≤ 0.05), which was undetectable at 72 h, which is in line with expression patterns seen in Figure 1A and Figure 1B.

### h*CFTR* (c)mRNA and hCFTR protein quantification in lungs after application *in vivo*

We tested for the localization of h*CFTR* (c)mRNA complexed with nanoparticle in the lungs after i.t. or i.v. application via RT-qPCR, quantified the hCFTR protein expression with hCFTR ELISA and then evaluated its immunogenicity depending on modification. Accordingly an experimental setup has been established (Figure 2A) with comprehensive treatment schemes and unambigous main outcome parameters (Figure 2B). In contrast to the *in vitro* data, when 40 μg 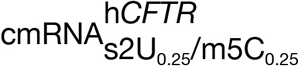 was i.v. injected into the mice, this resulted in a ~3.8-fold higher accumulation of that mRNA in the lung as compared to 40 μg 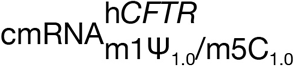 and h*CFTR* pDNA (*P* ≤ 0.05, Figure 2C). More importantly, we wanted to analyze if there is a significant increase in hCFTR protein levels in the lungs of treated mice by hCFTR ELISA (Figure 2D). These analyses confirmed that mice treated with 40 μg 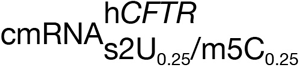 i.v. had a highly significant increase of hCFTR in the lungs of treated mice vs control mice (*P* ≤ 0.01; Figure 2D). Moreover, we tested the effects of an increased amount of 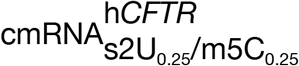 and h*CFTR* pDNA i.t. to 80 μg, which initially seemed to have a low deposition (Figure 2D), but again showed a clear and significant increase of hCFTR protein compared to control mice (Figure 2D) (*P* ≤ 0.05). All the mock controls used in hCFTR Elisa has proved to be not significantly different from negative control.

**Fig. 2:**
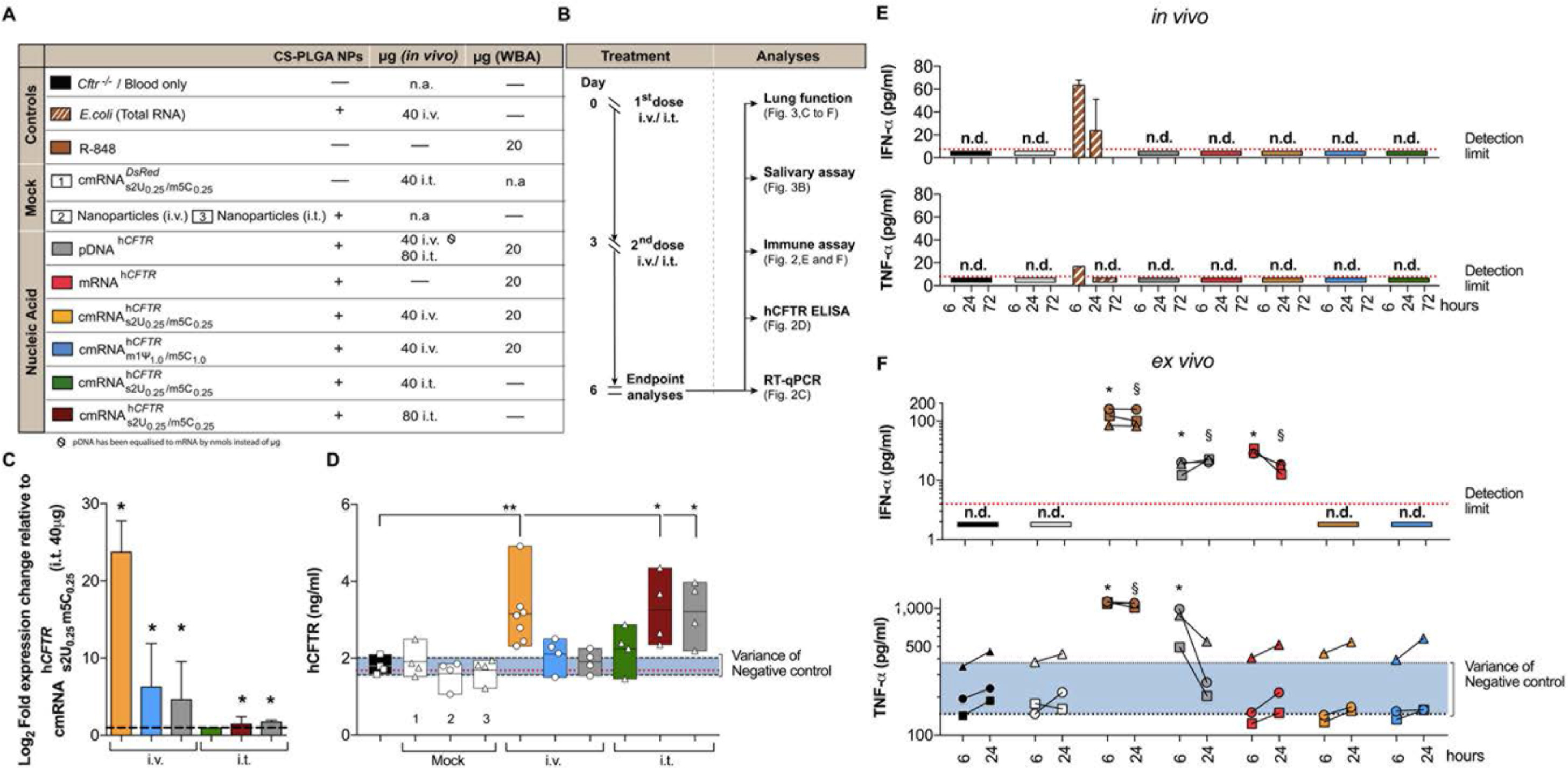
*In vivo* study plan, expression of modified h*CFTR* mRNA and hCFTR protein in mouse lungs and immunogenicity in mice and human whole blood. (**A**) All mouse groups, particles and particle combinations depicted in the study plan (**B**) and utilized in (**C-F**) are color-coded for their treatment schemes, including dosage and application routes. (**C**) Relative amounts of differently modified h*CFTR* mRNAs in the lungs, applied i.v. or i. t., then determined by RT-quantitative PCR, compared to 40μg 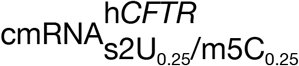., i.t. injection (⋆, *P* ≤ 0.05); *n* = 4-7 mice per group. (**D**) ELISA, detecting specifically human CFTR, was performed on lung preparations at day 6 (endpoint); the same *n* = 4-7 mice per group as in (**C**) were used. ⋆, *P* ≤ 0.05, ⋆⋆, *P* ≤ 0.01 versus untreated *CFTR* knock-out mice. (**E**) Mice were i.v. or i.t. injected with a mix of (c)mRNA and NPs at a 1:10 ratio, and ELISAs were performed post-i.v./i.t.-injection at three different time points. n.d., not detectable. (**F**) 2 pl whole blood, each from three different healthy human donors, were incubated with either R848 (1 mg/ml) or 3.82 pMol pDNA or 7.91 pMol (c)mRNA (providing the same total number of nucleic acid molecules) and NPs at a 1:10 ratio; after 6 h and 24 h the immune response was determined by ELISA in the sera; ⋆ and §, *P* ≤ 0.05 versus control at 6 h and 24 h, respectively. The red dotted lines in (**D-F**) mark the detection limit as specified in the respective ELISA kit. The blue areas in (**D, F**) represent the variance of the negative controls which are biological replicates. n.d., not detectable. All bar graph data are depicted as the means ± SD and box plots data are represented as the means ± minimum to maximum values.

### h*CFTR* (c)mRNA immunogenicity *in vivo* in mice after i.v. application and *ex vivo* in an adapted human whole blood assay⋆

⋆all *in vivo* experiments were performed with nanoparticles if not stated otherwise Due to lack of a reliable method to detect immune responses that therapeutic mRNAs may trigger in a living orgamisms, we focussed on two different approaches. First, we applied different compounds such as nanoparticles and R-848 (Resiquimod, a strong TLR7 and TLR8 agonist) and modified or unmodified mRNA i.v. or i.t. to mice to monitor their immune reaction at three different time points. All compounds, mRNAs and application routes are color-coded in Figure 2A. Surprisingly, applying 40 μg unmodified h*CFTR* mRNA or h*CFTR* cmRNA (with any modifications used) did not lead to detectable responses of key cytokines IFN-α or TNF-α (detected by ELISA) at all three time points (Figure 2E) (28, 29). Nanoparticles alone (used in all *in vivo* experiments) showed no immune response over the detection limit. However, as expected the positive control (*E. coli* extract total RNA) resulted in a significant increase of IFN-α and TNF-α at 6 h and a trend increase of IFN-α at 24 h, while an effect at 72 h was not detectable (Figure 2E).

In contrast to that, different results were obtained when we used a more complex assay based on human whole blood. Interestingly, the negative control groups (blood only and NP only) did not raise IFN-α values above the detection limit (Figure 2F, red dotted lines), while one sample of TNF-α was already measureable in human blood untreated or treated only with NPs. That is the reason why we adapted the graphical presentation of Figure 2F as we already did in Fig. 2D, using a blue colored area that represents the variance of the negative controls, which are biological replicates. The positive control (R-848) lead to a strong and significant production of both IFN-α (6 h and 24 h, respectively; *P* ≤ 0.05) and TNF-α (6 h and 24 h, respectively; *P* ≤ 0.05) (Figure 2F).

Human whole blood transfected with h*CFTR* cmRNAs showed a very similar result in cytokine expression as observed for negative controls: the IFN-α levels did not reach the detection limit of the ELISA; TNF-α responses were not statistically significant at 6 h and 24 h, respectively (Figure 2F). unmodified h*CFTR* mRNA resulted in a significant increase of IFN-α (30.6 ± 3.0 pg/ml and 16.6 ± 3.5 pg/ml at 6 h and 24 h, respectively; *P* ≤ 0.05) while the TNF-α levels were in line with the negative control. While h*CFTR* pDNA triggered high TNF-α responses at 6 h and 24 h (785.5 ± 256.80 pg/ml and 336.29 ± 182.68 pg/ml respectively; *P* ≤ 0.05), lower but detectable IFN-α responses after 6 h and 24 h (17.24 ± 4.43 pg/ml and 21.82 ± 1.21 pg/ml) could be observed. Due to both a significantly lower expression of unmodified h*CFTR* mRNA *in vitro* (Figure 1A and 1B) and higher immune responses of unmodified h*CFTR* mRNA depicted in Figure 2F, we focused on h*CFTR* cmRNAs and h*CFTR* pDNA in the following therapeutic studies.

### Therapeutic effect of h*CFTR* (c)mRNA *in vivo* in mice after i.t. and i.v. application⋆

⋆all *in vivo* experiments were performed with nanoparticles if not stated otherwise After the expression- and immuno-profiling, we investigated the therapeutic potential of cmRNA in a mouse model of Cystic Fibrosis. In order to test the efficacy of h*CFTR* cmRNA, *CFTR* knock-out mice have been used in several experimental settings that are explained and color-coded in Figure 3A. First, we performed a well established functional test, measuring the mouse saliva chloride concentration (30). The saliva chloride concentration detected in *Cftr^-/-^* mice (4084 ± 236.8 ng/μl) was significantly higher compared to *Cftr*^+/+^ mice (748.8 ± 96.9 ng/μl, *P* ≤ 0.01; Figure 3B). The treatment with either 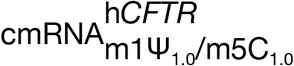 i.v. or i.t. (80μg) significantly lowered the chloride concentrations in the saliva of *Cftr*^-/-^ mice more than 52% and 36% repectively (*P* ≤ 0.05; Figure 3B). However 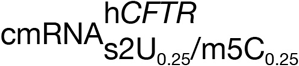 and h*CFTR* pDNA treated mice (i.v.) only provided a 22% reduction, although increased amount h*CFTR* pDNA treatment (i.t.) resulted in 30% reduction of chloride concentration in saliva of *Cftr*^-/-^ mice (*P* ≤ 0.05; Figure 4B).

**Fig. 3:**
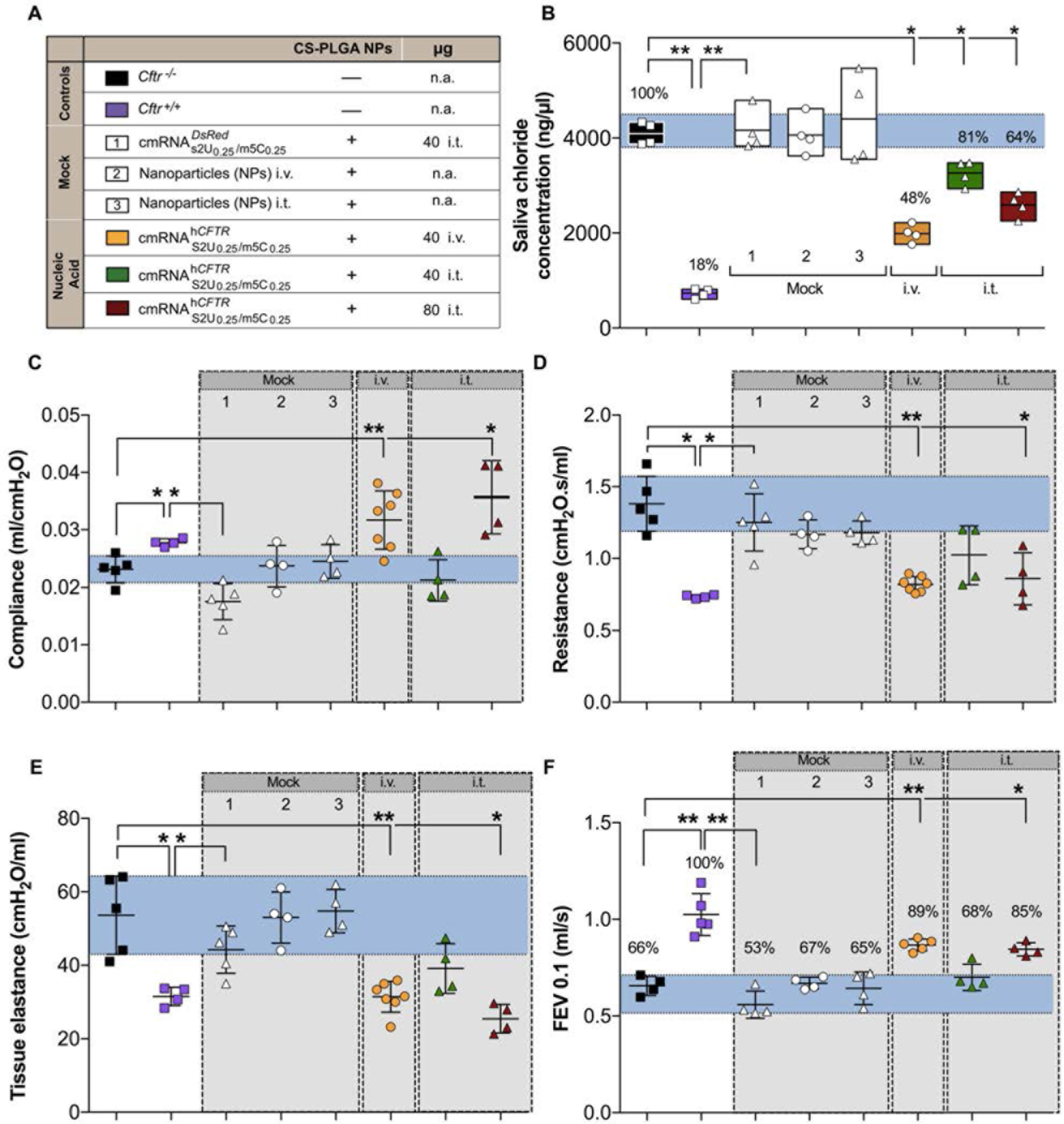
*In vivo* lung function measurements in h*CFTR* mRNA treated *CFTR* knock-out mice. All mouse groups utilized in (**B-F**) are color-coded for their treatment schemes (**A**), including dosage and application routes. **B**) Functional test of reconstituted CFTR channel compared to *Cftr* knock-out mice (black), positive controls (violet), and percentages relative to the positive control; *n* = 4-7 mice per group; 3 mock controls were included (white); boxes represent the means ± minimum and maximum values. (**C-F**) Precision *in vivo* lung function measurements covering all relevant outcome parameters on mice treated twice (see **A**) and measured 72 hours after the 2^nd^ installment; *n* = 4-7 mice per group. Data represent the means ± SD on compliance, resistance, tissue elastance and Forced Expiratory Volume in 0.1 seconds (FEV_0.1_). ⋆, *P* ≤ 0.05, ⋆⋆, *P* ≤ 0.01 versus untreated *Cftr* knock-out mice.

To assess the impact of h*CFTR* cmRNA on lung function, we evaluated clinically relevant parameters using the FlexiVent^®^ lung function measurement system. We observed significant differences between Mock controls / *Cftr*^-/-^ and healthy wild-type mice for all parameters measured (*P* ≤ 0.05; Figure 3C-E and *P* ≤ 0.01; Figure 3F). Applying 40 μg of 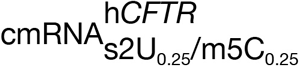 i.v. significantly lowered the resistance (*P* ≤ 0. 01; Figure 3D). Furthermore, i.v. administration of 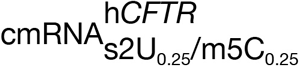 significantly increased the compliance from 0.02 ± 0.01 ml/cmH_2_O (*Cftr*^-/-^ mice) to 0.03 ± 0.01 μl/cmH_2_O (*P* ≤ 0.01), reaching equivalent values to those measured in *Cftr*^+/+^ mice (Figure 3C). In the i.t. treated groups, a significant improvement of resistance and compliance could be detected when the 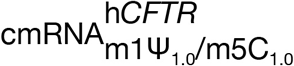 dose was increased to 80 μg (0.86 ± 0.18 cmH_2_O.s/ml and 0.04 ± 0.01 ml/cmH_2_O, respectively; *P* ≤ 0.05; Figures 3C-D). However 40 μg of 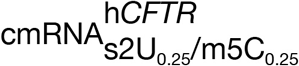 i.v. lowered the resistance but not as effectively as 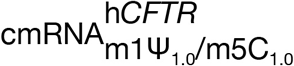 (*P* ≤ 0.05; Figure 4D) and h*CFTR* pDNA (80μg) i.t. treated mice also produced significant improvement of resistance and compliance (*P* ≤ 0.05, Figure 4C-D). Furthermore the i.v. application of 40 μg 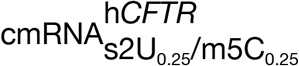 or h*CFTR* pDNA did not alter compliance significantly (Figure 4C).

**Fig. 4:**
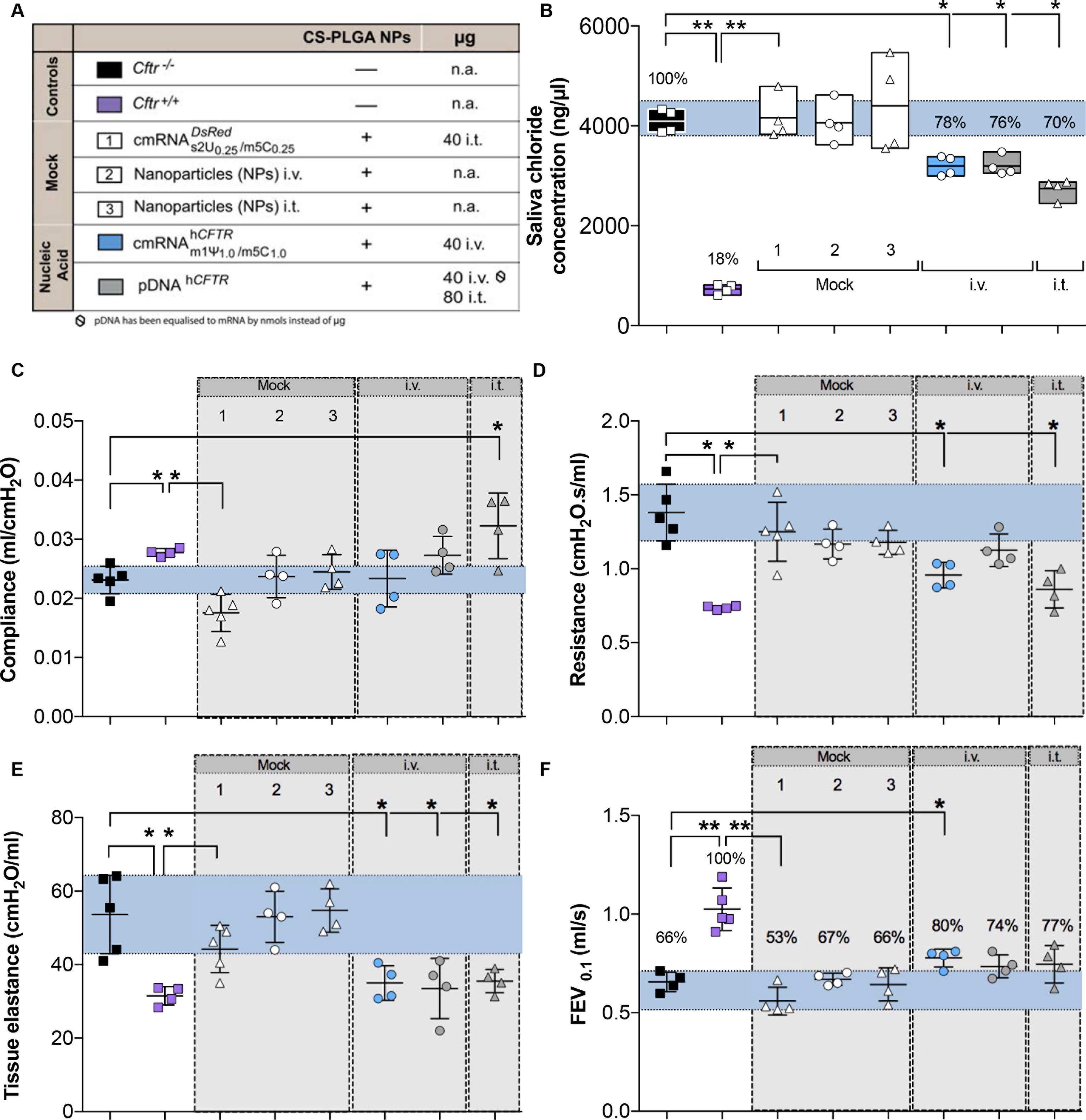
*In vivo* lung function measurements in h*CFTR* pDNA treated *CFTR* knock-out mice. All mouse groups utilized in (**B-F**) are color-coded for their treatment schemes (**A**), including dosage and application routes. **B**) Functional test of reconstituted CFTR channel compared to *Cftr* knock-out mice (black), positive controls (violet), and percentages relative to the positive control; *n* = 4-7 mice per group; 3 mock controls were included (white); boxes represent the means ± minimum and maximum values. (**C-F**) Precision *in vivo* lung function measurements covering all relevant outcome parameters on mice treated twice (see **A**) and measured 72 hours after the 2^nd^ installment; *n* = 4-7 mice per group. Data represent the means ± SD on compliance, resistance, tissue elastance and Forced Expiratory Volume in 0.1 seconds (FEV_0.1_). ⋆, *P* ≤ 0.05, ⋆⋆, *P* ≤ 0.01 versus untreated *Cftr* knock-out mice.

Tissue elastance (peripheral lung mechanics), that is, energy conservation in the alveoli, was significantly improved for 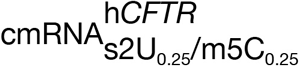 i.v. treated group (*P* ≤ 0.01; Figure 3F) and i.t. application of 80 μg 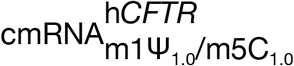 (*P* ≤ 0.05; Figure 3E) compared to *Cftr*^-/-^ mice, in which tissue elastance was detected with a value of 53.61 ± 10.67 cmH_2_O/ml. 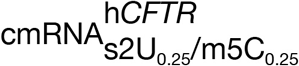 and h*CFTR* pDNA, (i.v. and i.t.) treatment produced a improved tissue elsastance but failed to reach the effect of i.v treatement of 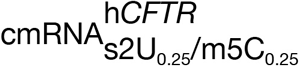 (*P* ≤ 0.05; Figure 4E and Figure 3E).

FEV_0.1_ (human equivalent of FEV_1_) of *Cftr*^+/+^ mice defined as projecting 100% forced exhale volume, the i.v. injection of 40 μg 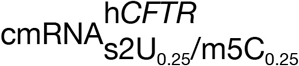 or i.t. application of 80 μg 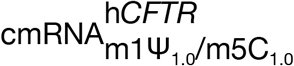 improved the FEV_0.1_ by 23% (*P* ≤ 0.01) and 19% (*P* ≤ 0.05) respectively compare to untreated *Cftr*^-/-^ (Figure 3F). Only i.v. injection of 40 μg 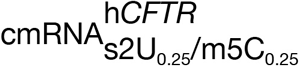 provide a FEV_0.1_ improvement of 14% which is satistically significant (*P* ≤ 0.05; Figure 4F). However i.v. or i.t. administration of h*CFTR* pDNA showed no significant improvement of FEV_0.1_. Taken together, these results demonstrate significant lung function improvement in all relevant lung function parameters of *Cftr*^-/-^ mice treated with h*CFTR* cmRNA.

## Discussion

Although much progress has been achieved since the discovery of the *CFTR* gene 25 years ago, there is still a substantial need to restore robust CFTR function in patients suffering from cystic fibrosis (31). With the recent approvals of the small molecule agents ivacaftor and lumacaftor, science has paved a possible way to overcome the hurdles caused by the disease-conferring gene. Those treatments can be more or less effectively applied to patients bearing *CFTR* mutations delF508 (Lumacaftor- ivacaftor/Orkambi) and G551D (ivacaftor) (32–35). However, lung function as one of the main outcome parameters probably having the most significant influence on life quality of CF patients, is rarely tested in preclinical models. In fact, actual effects of (modern) existing drugs on lung function, with forced expiratory volume in one second (FEV_1_) as a key parameter, are quite low (36). Here by using h*CFTR* (c)mRNA, we are presenting a proof of concept for a viable and potent therapeutic alternative. We have vigorously tested mRNA therapy with focus on *in vivo* lung function normalization while avoiding any possible, unwanted immune reponses for a possibility of repeated dosing. The unique formulation utilized, can be used both topically (intratracheally) and systemically (via i.v. injection), having in both cases a profound effect on normalizing the lung function parameters, including compliance, resistance and FEV_01_ of treated *Cftr*^-/-^ mice to values obtained from *Cftr*^+/+^mice.

*In vitro*, using h*CFTR* cmRNA, CFTR protein expression in CFBE cells was increased up to 5.5-fold compared to unmodified h*CFTR* mRNA, which is consistent with previous studies obtained by us and others (2, 26, 37). Incorporation of naturally occurring nucleosides has been shown to suppress inhibitory effects on translation by avoiding detection by pattern recognition receptors (PRRs) such as Toll-like receptors (TLRs) TLR3, TLR7 and TLR8 (28, 29). Those receptors play a crucial role in the detection, processing and degradation of mRNA. Interestingly, depending on the mRNA modification, kinetics of hCFTR expression varies upon the different nucleosides used. In fact, we did not observe an increased quenching efficacy after 72h in CFTR null A549 cells, which would corroborate our findings from western blot analyses. Although there is a significant increase in I^−^ influx by functional hCFTR channels at 48h post transfection, both modified hCFTR mRNAs showed similar activity. Consequently, we assume that upon different cell lines, kinetics by which the hCFTR protein is expressed varies. Earlier studies support our notion that different modified mRNAs can have an impact on the translational effect between distinct cell lines (26, 29).

To better determine the clinical potential of CFTR-encoded cmRNA we compared not only different modifications *in vivo* but also two different routes of administration. Applying cmRNA i.t., has been shown to significantly prolong survival in a surfactant protein-B mouse model (22). Given the fact that in patients suffering from CF one of the key barriers is the airway mucus layer in which inhaled particles are more likely to get trapped and removed, we sought to apply h*CFTR* cmRNA/pDNA complexed to NPs by i. v. injection as an alternative administration route. Systemic delivery via lipid modified polymeric nanoparticles has been already shown to target the lungs efficiently (38). In this study, by applying h*CFTR* cmRNA consecutively, both modifications were successfully delivered to the lungs with the i.v. route being more efficient at doses of 40 μg (2 mg/kg) per treatment. Intriguingly, in contrast to the results obtained *in vitro*, 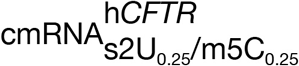 showed a significantly higher CFTR protein expression with higher accumulation of h*CFTR* cmRNA in lung cells. Assuming differences of cmRNA- encoded transgene expression between distinct cell lines, it is plausible to consider such differences between *in vitro* versus *in vivo* applications, which is by far more complex. In this respect, the higher amount of cmRNAh*CFTR*,s2U_0.25_/m5C_0.25_ found in lung cells after i.v. injection, might be due to the fact that its nucleoside composition is more favorable to evade PRRs, thus being less degraded. However, regardless of cmRNA kinetics we also observed differences in the delivery route of h*CFTR* cmRNAs/pDNA-NPs. Our data suggests i.v. injection to be more efficient in delivering such complexes to the lung than topical administration. Tests of h*CFTR* cmRNA-NP’s capacity of mucus penetration are in planning phase. The upper airways are lined with mucus and mucociliary movements clear foreign particles immediately. In addition, the main barriers in the deeper areas are the alveolar lining, scavenger transporters and alveolar macrophages (39, 40). We therefore concluded, that the dosing by which cmRNA-NPs were delivered i.t. was not sufficient to reach the lung cells efficiently. Indeed, increasing the amount by doubling the dose (to 80 μg) for each treatment showed a significant increase in hCFTR expression.

To exclude immune reactions caused by either NPs or the (c)mRNA itself, we conducted extensive immune assay tests *in vivo*. Except for the positive control (*E. coli* total mRNA) we could not detect any immunostimulatory effect *in vivo* that could arise from NPs or the (un-)modified h*CFTR* mRNAs. These results confirm our previous studies in which we showed that NPs as well as modified mRNA could be administered safely to the lungs without any substantial increase in cytokines, or inflammatory-related cells such as macrophages or neutrophils (22). Systemic delivery has also been reported to have no impact on proinflammatory cytokine secretion (24). To better mimic the *in vivo* human conditions, we performed an *ex vivo* whole blood assay (WBA) which offers a more complex environment to test for immune responses. This assay has already been used in a number of preclinical settings and Coch and colleagues could demonstrate that it has the potential to reflect broad aspects of the *in vivo* cytokine release caused by oligonucleotides (41). Indeed, we could show that the small molecule resiquimod (serving as a positive control by activating TLR7 and TLR8) lead to a substantial release of IFN-α and TNF-α. plasmid-encoded hCFTR as well as unmodified h*CFTR* mRNA also showed elevated cytokine levels probably due to the activation of innate immune receptors (28, 29). In contrast, incorporation of modified nucleosides into h*CFTR* mRNA abolished such responses, with no detectable amounts of IFN-α. This is in concert with previously published data, demonstrating cmRNA’s limiting immune responses, mainly by evading detection from receptor such as TLRs, RIG-1, MDA-5 or PKR (28, 37). Interestingly, even though TNF-α could be detected, it rather shows donor-dependency than effects deriving from NPs and/or h*CFTR* cmRNA with cytokine levels being all within the variance of negative controls. Although it mirrors only the blood compartment and does not reflect the more complex *in vivo* situation, the WBA can give a prediction of how cytokines are released in the human system in response to systemically applied (c)mRNA prior to clinical testing.

Eventually, we determined the impact of h*CFTR* cmRNA and plasmid-encoded hCFTR on relevant physiological outcomes such as the saliva chloride concentration as well as important lung function parameters to evaluate its therapeutical effect. Sweat chloride concentration has become an accepted method as a diagnostic readout to assess treatment effects of CF patients (42). As an analog, chloride concentration of β- adrenergic stimulated salivary glands of CFTR knock-out mice can be investigated as it complies with findings in CF patients (30). In this study, we could show a substantial difference in salivary Cl” content of h*CFTR* cmRNA and h*CFTR* pDNA treated mice - both, i.v and i.t. - compared to their untreated counterpart. With end point-analysis, a significant decrease in Cl^−^ to nearly 60 % was observed, indicating a restoration of CFTR in the duct compartment of salivary glands and thus leading to an improved Cl^−^ absorption. Previous studies estimated that a restoration of CFTR activity to 50 % could lead to sweat chloride levels to near normal levels in CF patients. Given that, it is possible that h*CFTR* cmRNA treatment has the potential to improve CFTR activity to levels that are at least similar to those in patients with a mild CF phenotype (43).

To support our notion of improved CFTR activity, we additionally performed extensive lung function measurements using state-of-the-art technology to provide detailed *in vivo* information on different lung function parameters. *Cftr*^-/-^ mice have been criticized as a proper model for cystic fibrosis as it does not reflect the typical lung phenotype seen in CF patients (44). However, the reason behind that seems to be in how deeply lungs or other affected organs had been investigated. A layer of material can be observed with characteristics of an acid mucopolysaccharide on the bronchiolar surface and is also evident in alveoli by using scanning electron microscopy in *Cftr*^-/-^ mice, which is not evident in *Cftr*^+/+^ mice (45). Recent studies could clearly demonstrate reduced airway compliance and increased resistance in comparison to wild-type mice (46, 47). Indeed, we observed significantly higher and lower levels regarding resistance and compliance, respectively, in *Cftr*^-/-^ controls and mock-treated *Cftr*^-/-^ mice compared to homozygous wild type mice (*Cftr*^+/+^) mice and demonstrated that treatment with h*CFTR* cmRNA-NPs improved compliance and resistance significantly equal to those seen in healthy *Cftr*^+/+^ mice. FEV1 (Forced exhale volume) percentage (for mouse or small animal FEV _0.1_) is related to survival in CF and most important physiological parameter for CF patients. Previous study demonstrated that patients with a %FEV1 <30 had a 2-year mortality over 50% and hence it is regularly exhamined in clinical setup (48). Our study provide a significant improvement of FEV_0.1_ due to treatement with h*CFTR* cmRNA-NPs. Interestingly h*CFTR* pDNA when administrated via i.t. route improve other parameter of lung function measurements did amend FEV_0.1_ but not as significantly as h*CFTR* cmRNA-NPs.

In addition, intrinsic mechanical properties of the parenchyma are altered in *Cftr*^-/-^ mice with increased tissue elastance (47, 49). Such differences in peripheral lung mechanics indicate an energy loss due to frictional, resistive forces and that more work will be required to expand the lungs (elastance) (49). As for the central lung mechanics, peripheral measurements confirmed i.v.-treated groups to be more effective in targeting lung cells than topical application, thereby decreasing tissue elastance. We also observed i.t. administration of 80 μg 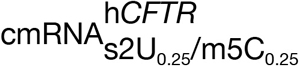 to positively compensate most of lung fuction parameters. Overall, we could demonstrate that certain protocols, applying h*CFTR* cmRNA either i.v. or i.t. efficiently restored lung function values equal to those of wild type. Suggesting a more evenly distribution through arteries and the bronchial circulation by i.v.-injection, especially for newborns and young infants, this route and formulation could lead to a very potent therapy. By providing functional CFTR early in life, the lungs could be protected from irreversable damage. Nevertheless, when applied intratracheally - which mimicks deep inhalation of a spray or powder formulation usually the primary application route in adults - an adjustment in dose and/or formulation (e.g. 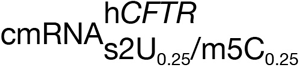 increased to 80 μg) might easily abrogate any negative effect of the *Cftr*^-/-^ genetic background on lung function.

Taken together, this study is the first proof of concept of efficient application of h*CFTR* cmRNA NPs *in vivo* to restore lung function in a *Cftr*-deficient mouse model. Importantly, we could neither detect immune responses *in vivo* nor in a more defined setting *ex vivo*. Applying h*CFTR* cmRNA to *Cftr* knock-out mice could efficiently restore lung function to levels of healthy control mice. In addition, our study compared - apart from two well-known mRNA modifications and h*CFTR* pDNA - also two different delivery routes, demonstrating that systemic administration of cmRNA targets lung cells more efficiently at lower dosages. This study provides a proof of concept for alternative treatment of patients suffering from CF. h*CFTR* cmRNA transcript supplementation may be broadly applicable for most *CFTR* mutations, not only in adults but already in the postnatal state, thereby protecting the lungs from exacerbations from the very beginning of life.

## Acknowledgements

We thank Dr. Dominik Hartl for experimental guidance on CFTR KO mice; Dr. Sandra Beer-Hammer and Dr. Franz Iglauer for helping to draft the respective animal proposal; Dr. Joachim Riethmüller for the numerous and fruitful discussions on translation of h*CFTR* mRNA therapy into the clinic, and Katrin Ganzenberg for mRNA isolation from various organs of *Cftr* mice and Brain weidensee for editing.

---

**Fig. S1:**
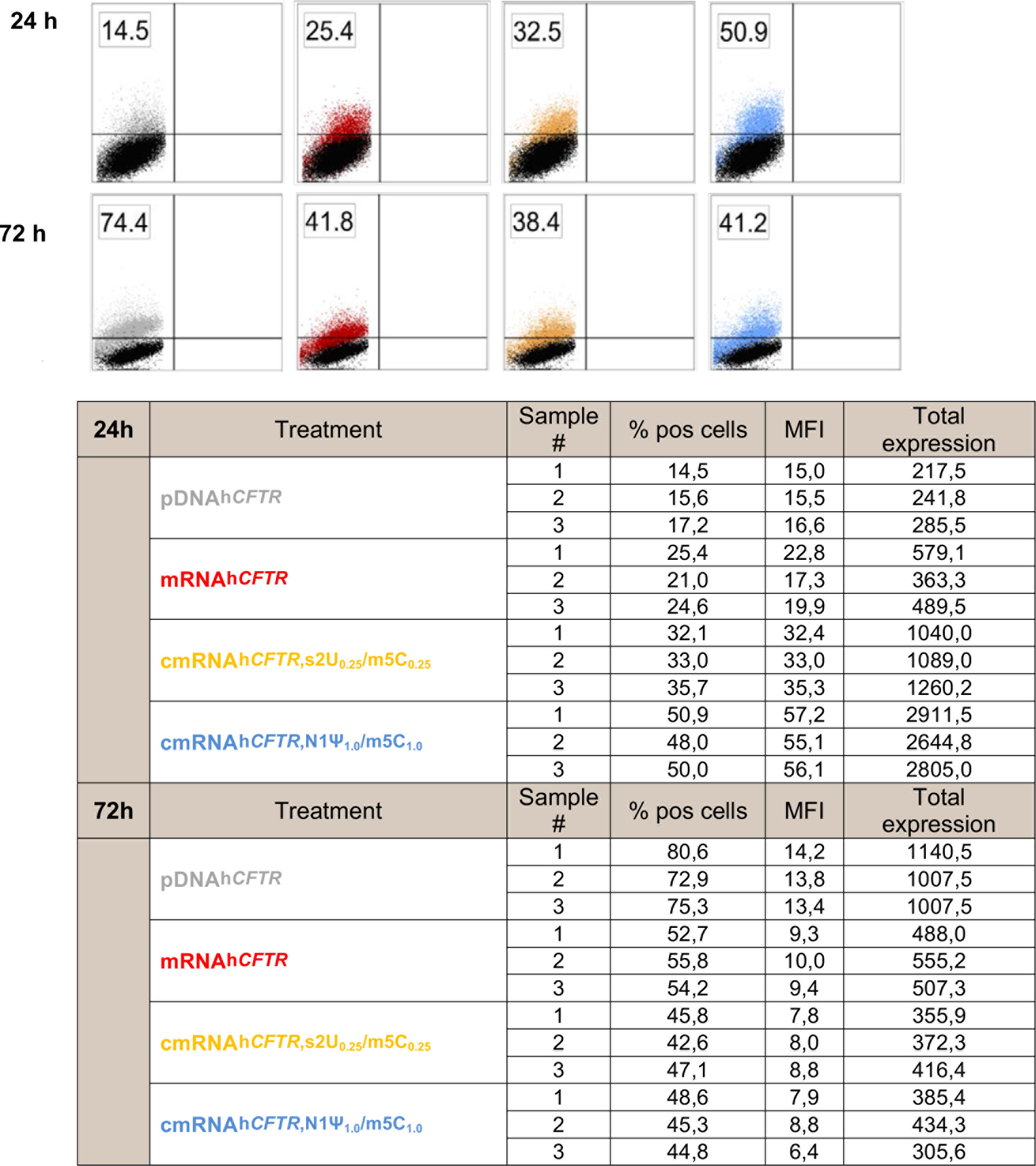
Expression analyses of hCFTR protein by flow cytometry; n=3

**Fig. S2:**
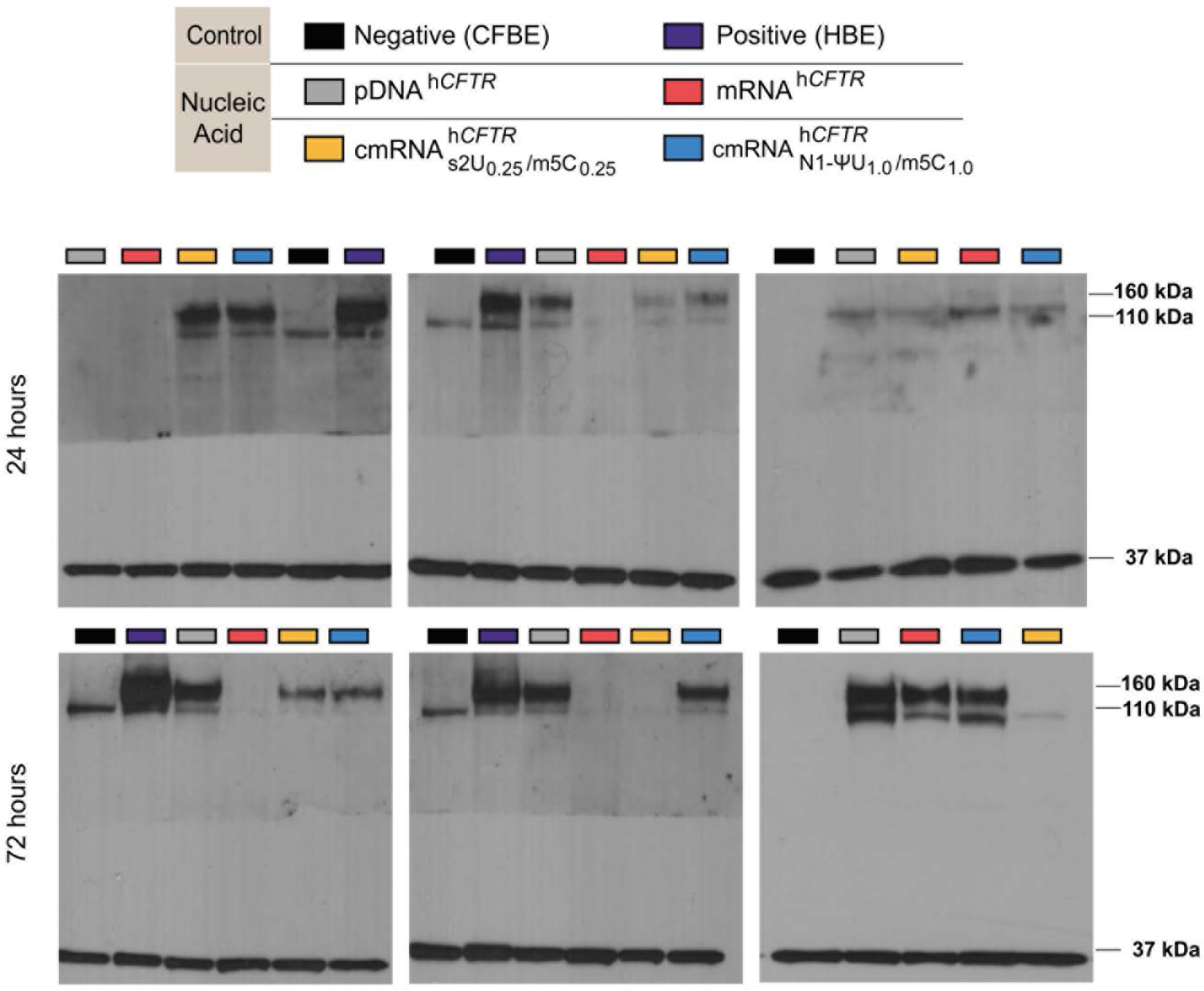
Western blot analyses of hCFTR protein in CFBE and HBE cells; n=3.

## Source data

**Fig. 1A:**
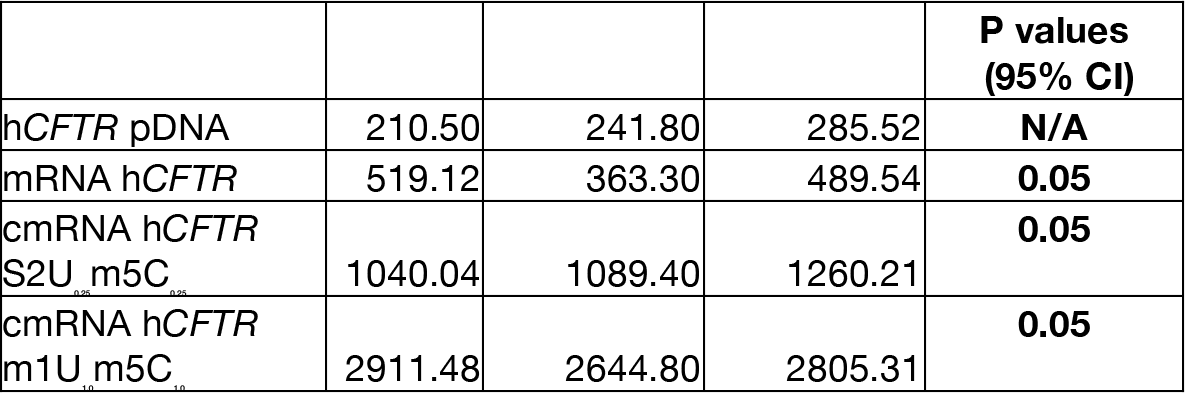
Total expression, **n=3**, 24hour- **Source 1** **Statistic:** Wilcoxon-Mann-Whitney test and *P* ≤ 0.05 was considered statistically significant

**Fig. 1A:**
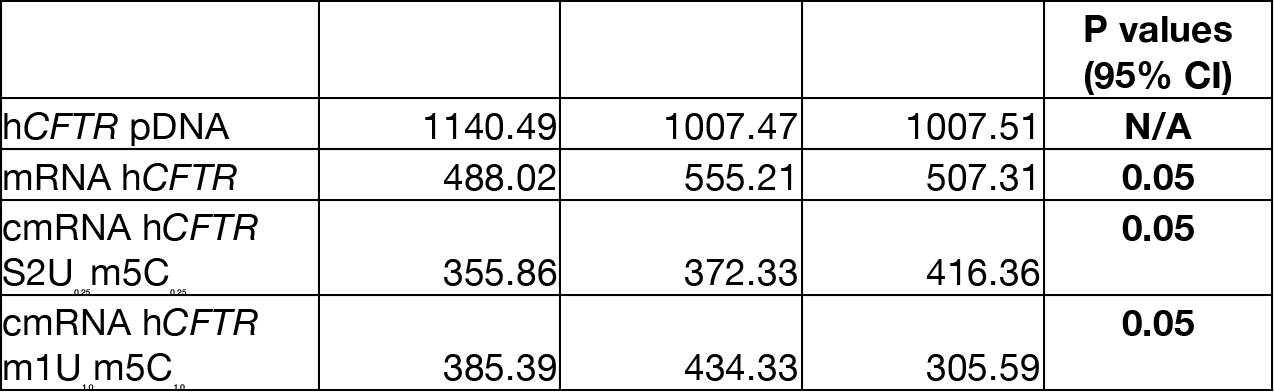
Total expression, **n=3**, 72hour- **Source 2** **Statistic:** Wilcoxon-Mann-Whitney test and *P* ≤ 0.05 was considered statistically significant

**Fig. 1A:**
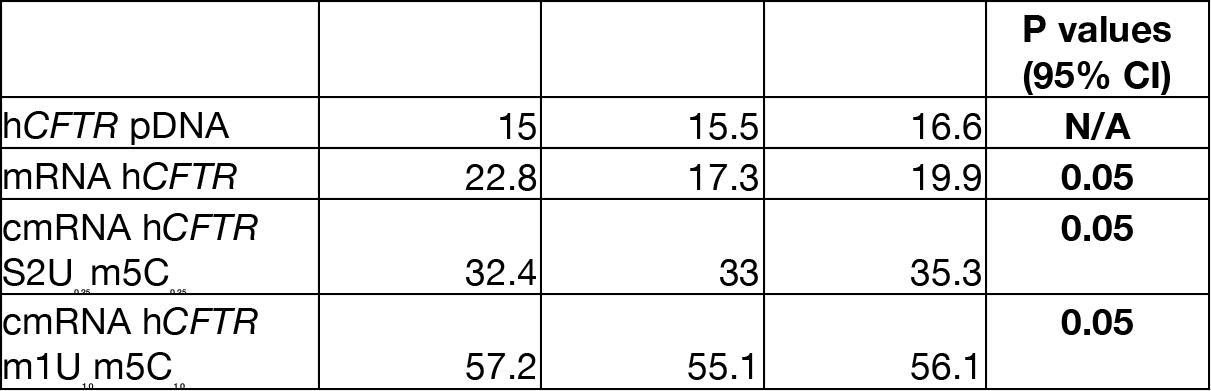
Median Fluorescence Intensity, **n=3**, 24hour- **Source 3** **Statistic:** Wilcoxon-Mann-Whitney test and *P* ≤ 0.05 was considered statistically significant

**Fig. 1A:**
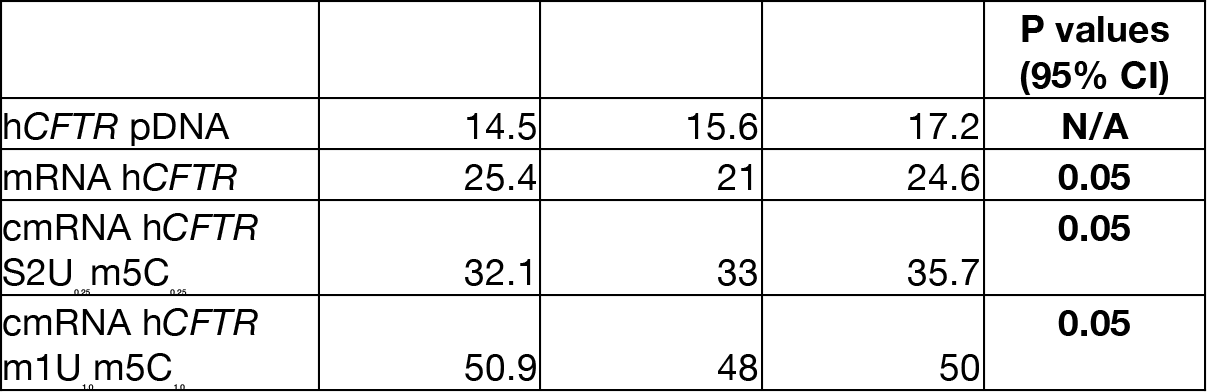
hCFTR Positive cell(%) **n=3**, 24hour- **Source 4** **Statistic:** Wilcoxon-Mann-Whitney test and *P* ≤ 0.05 was considered statistically significant

**Fig. 1A:**
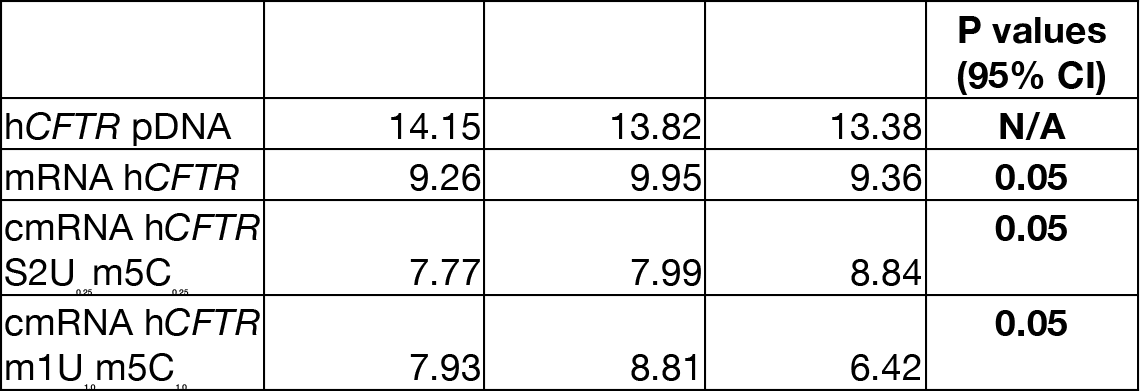
Median Fluorescence Intensity, **n=3**, 72hour- **Source 5** **Statistic:** Wilcoxon-Mann-Whitney test and *P* ≤ 0.05 was considered statistically significant

**Fig. 1A:**
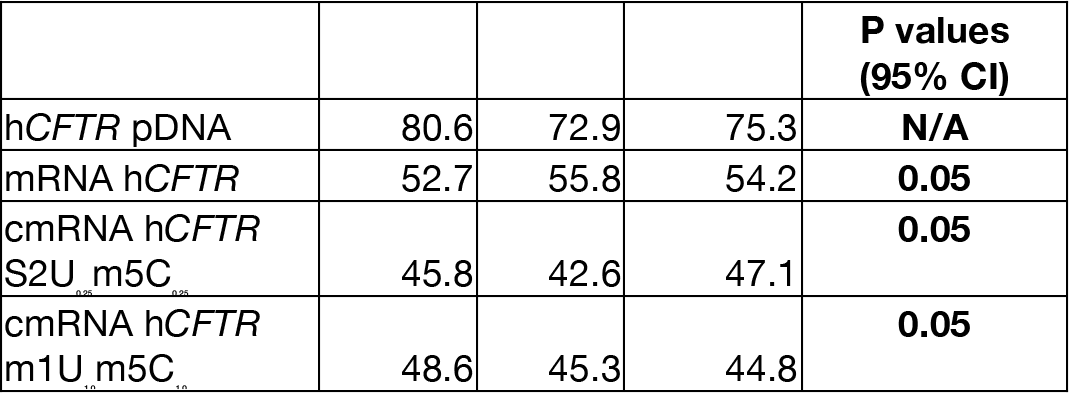
hCFTR Positive cell(%) **n=3**, 24hour- **Source 6** **Statistic:** Wilcoxon-Mann-Whitney test and *P* ≤ 0.05 was considered statistically significant

**Fig. 1B:**
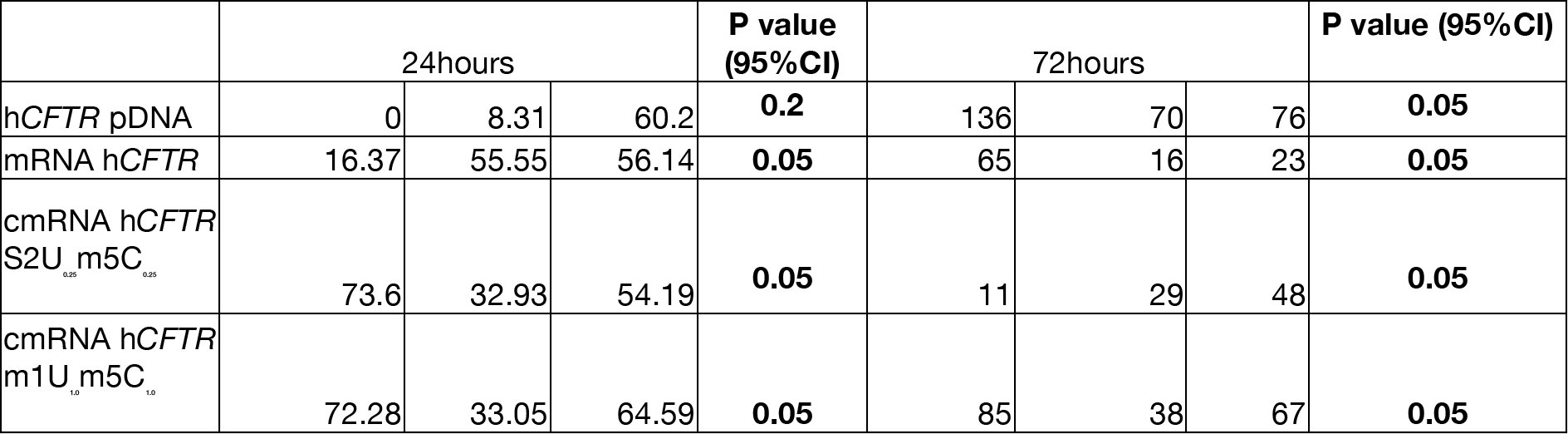
hCFTR expression by Western blot, n=3, Source 7 **Statistic:** Wilcoxon-Mann-Whitney test and *P* ≤ 0.05 was considered statistically significant

**Fig. 1C:**
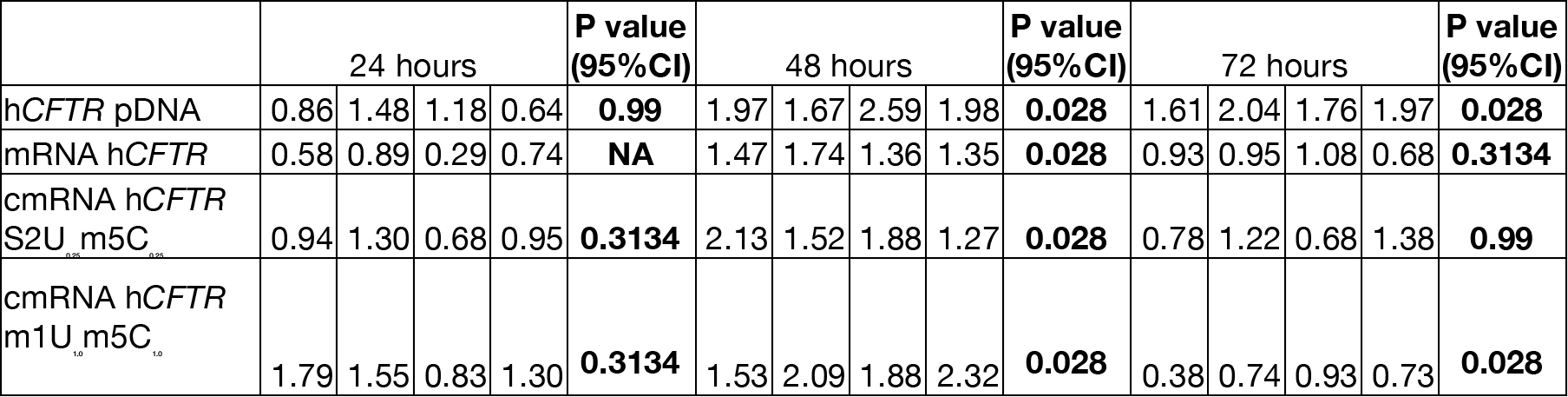
YFP assay for CFTR function **n=4**, for each time point. **Source 8** **Statistic:** Wilcoxon-Mann-Whitney test and *P* ≤ 0.05(two-sided) was considered statistically significant

**Fig. 2A:**
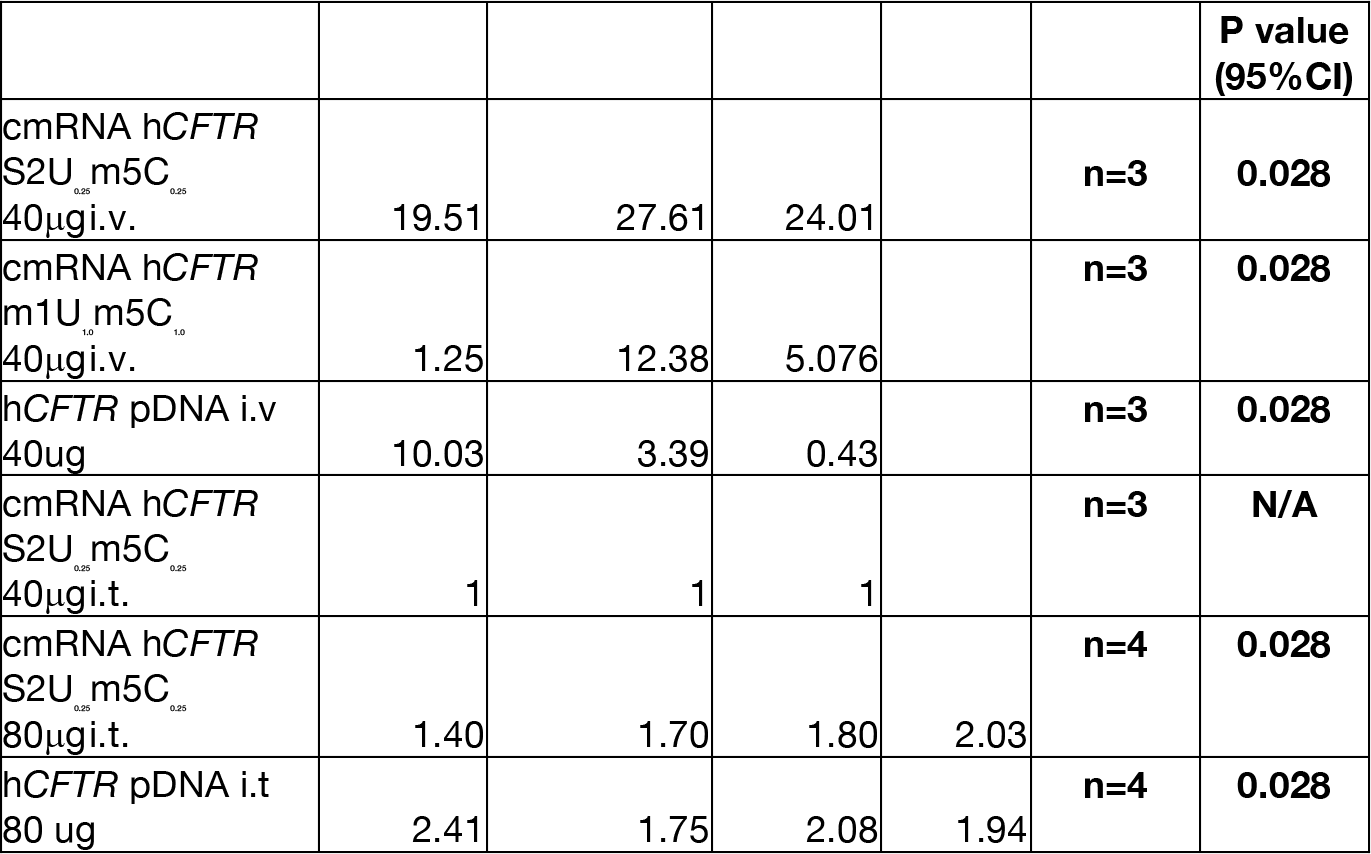
Expression fold change by qPCR. **Source 9** **Statistic:** Wilcoxon-Mann-Whitney test and *P* ≤ 0.05(two-sided) was considered statistically significant

**Fig. 2B:**
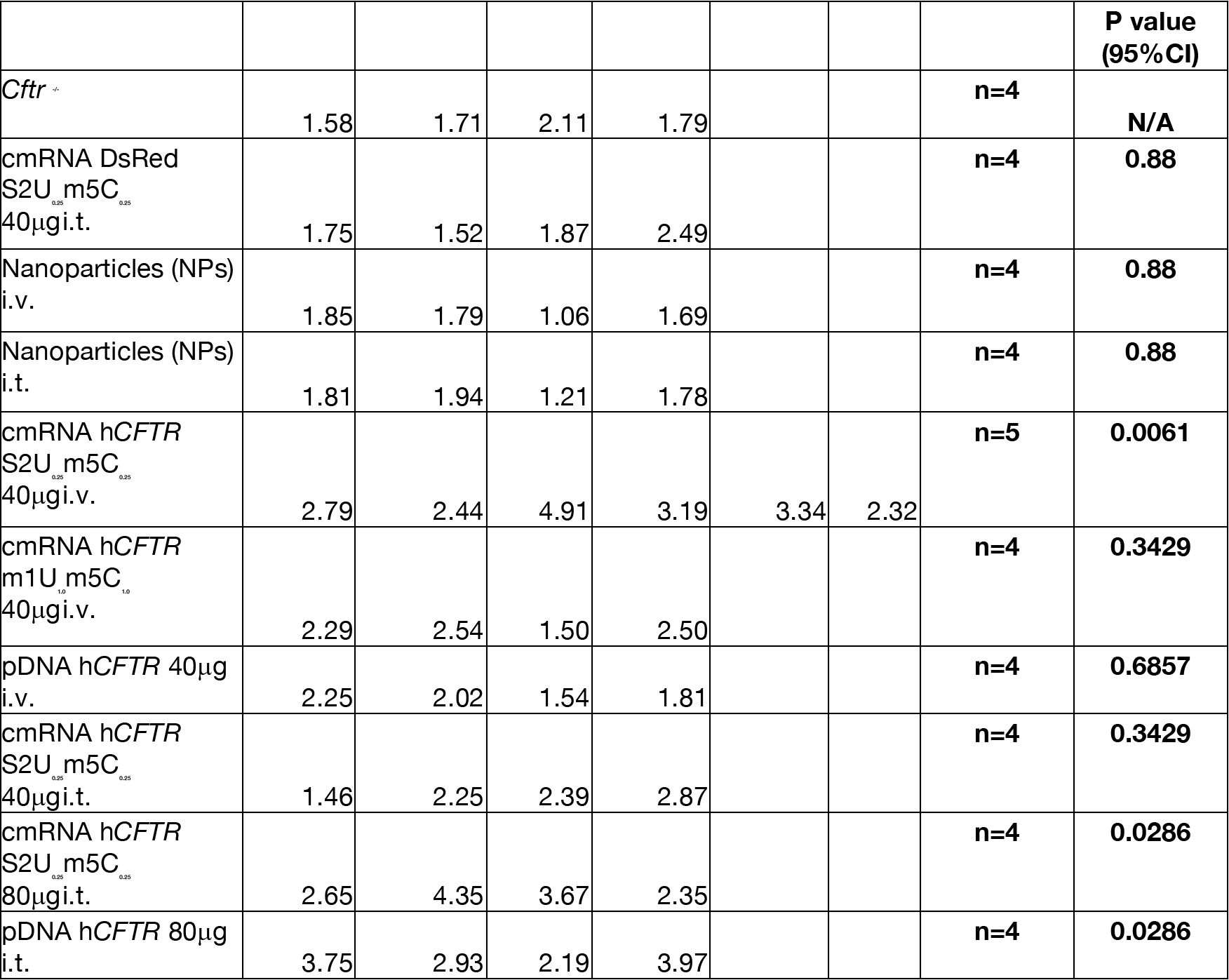
hCFTR Elisa. **Source 10** **Statistic:** Wilcoxon-Mann-Whitney test and *P* ≤ 0.05(two-sided) was considered statistically significant

**Fig. 2C:**
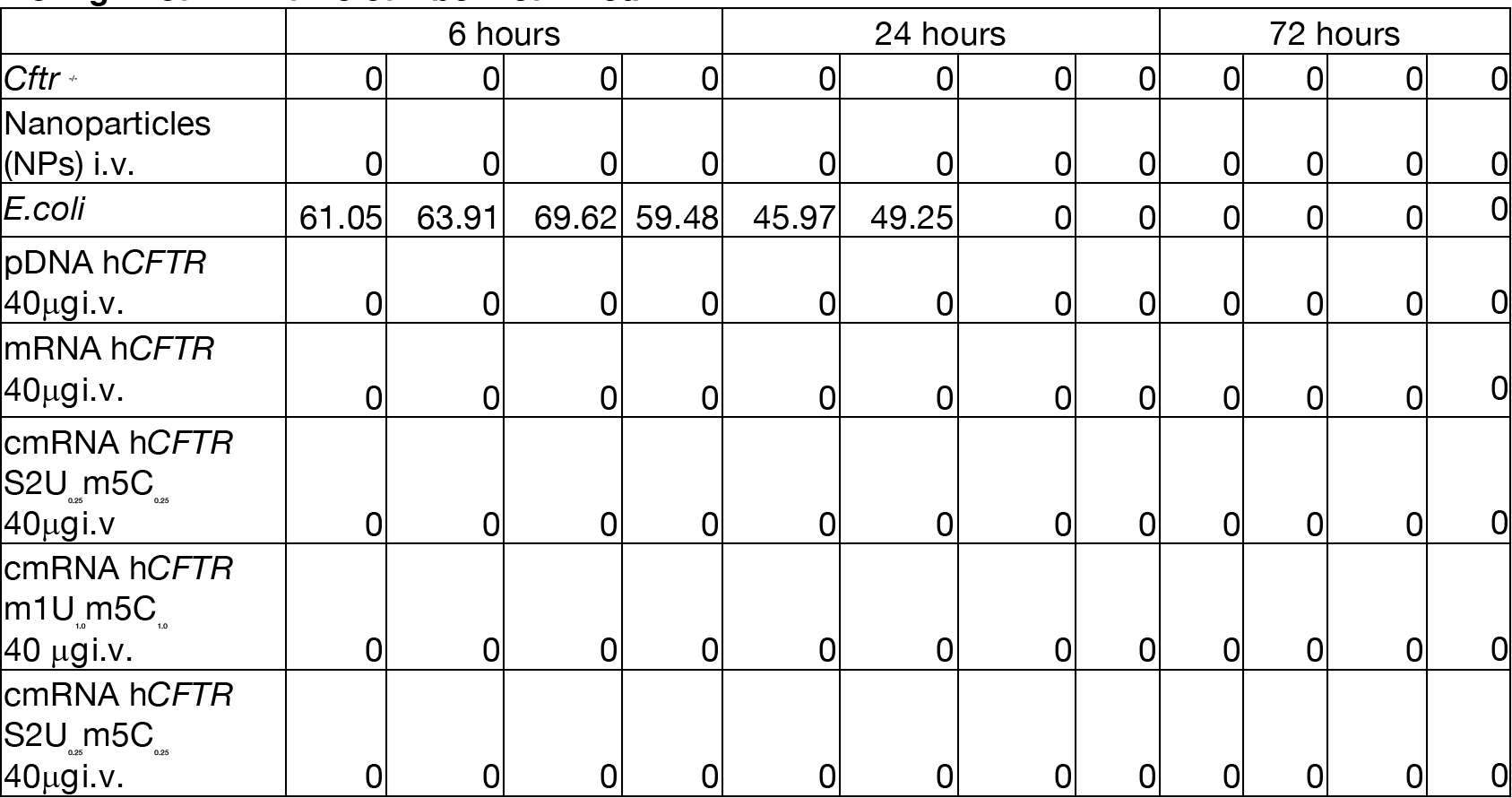
Immune Assay INF-Alpha *in vivo*, n=4 for each time point. Source 11. **Statistic:** Wilcoxon-Mann-Whitney test and *P* ≤ 0.05(two-sided) was considered statistically significant **No Significant P value can be measured**

**Fig. 2C:**
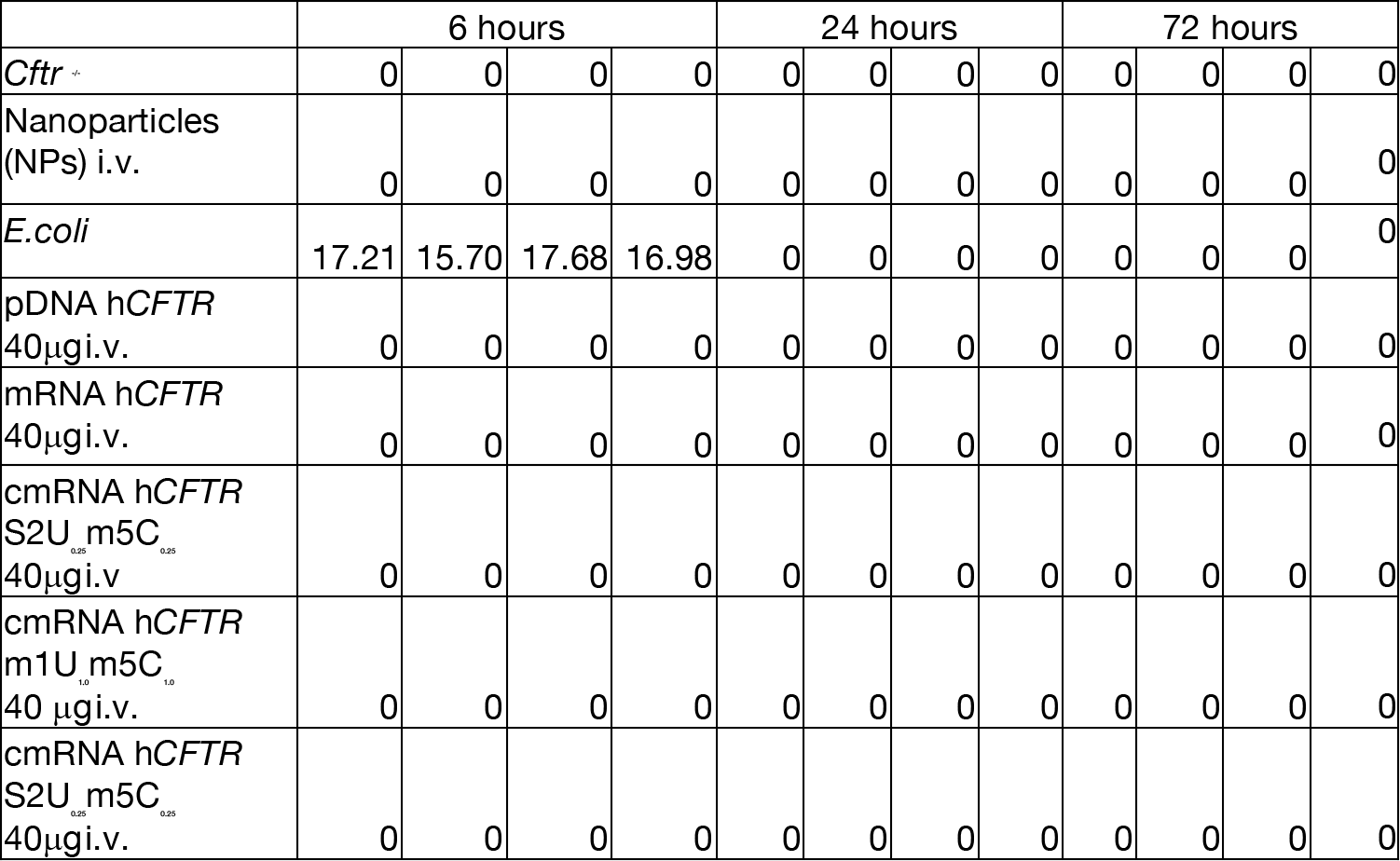
Immune Assay TNF-αlpha *in vivo*, n=4 for each time point. Source 12 **Statistic:** Wilcoxon-Mann-Whitney test and *P* ≤ 0.05(two-sided) was considered statistically significant **No Significant P value can be measured**

**Fig. 2D:**
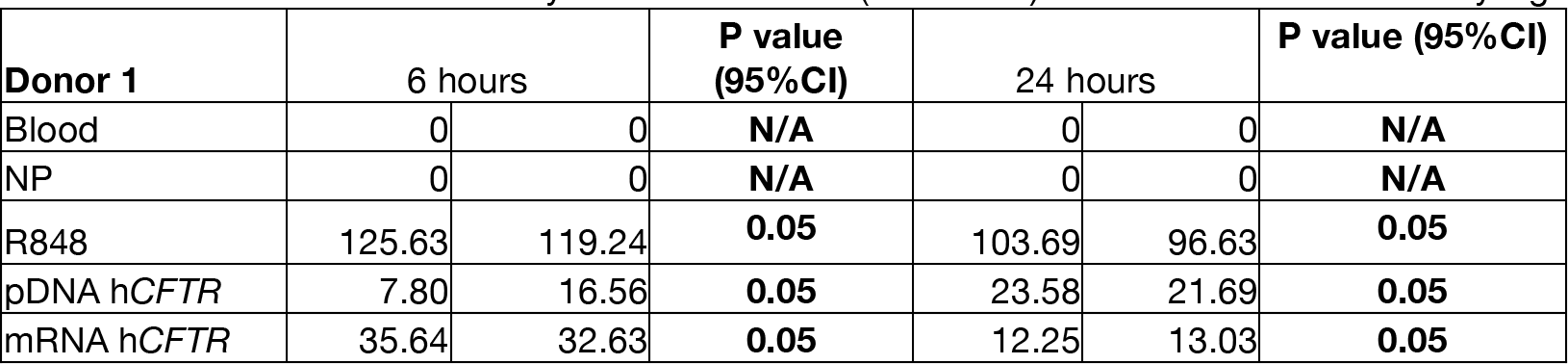
Whole blood assay INF-Alpha (*ex vivo*) Donor (n=3) Source 13 **Statistic:** Wilcoxon-Mann-Whitney test and *P* ≤ 0.05(two-sided) was considered statistically significant

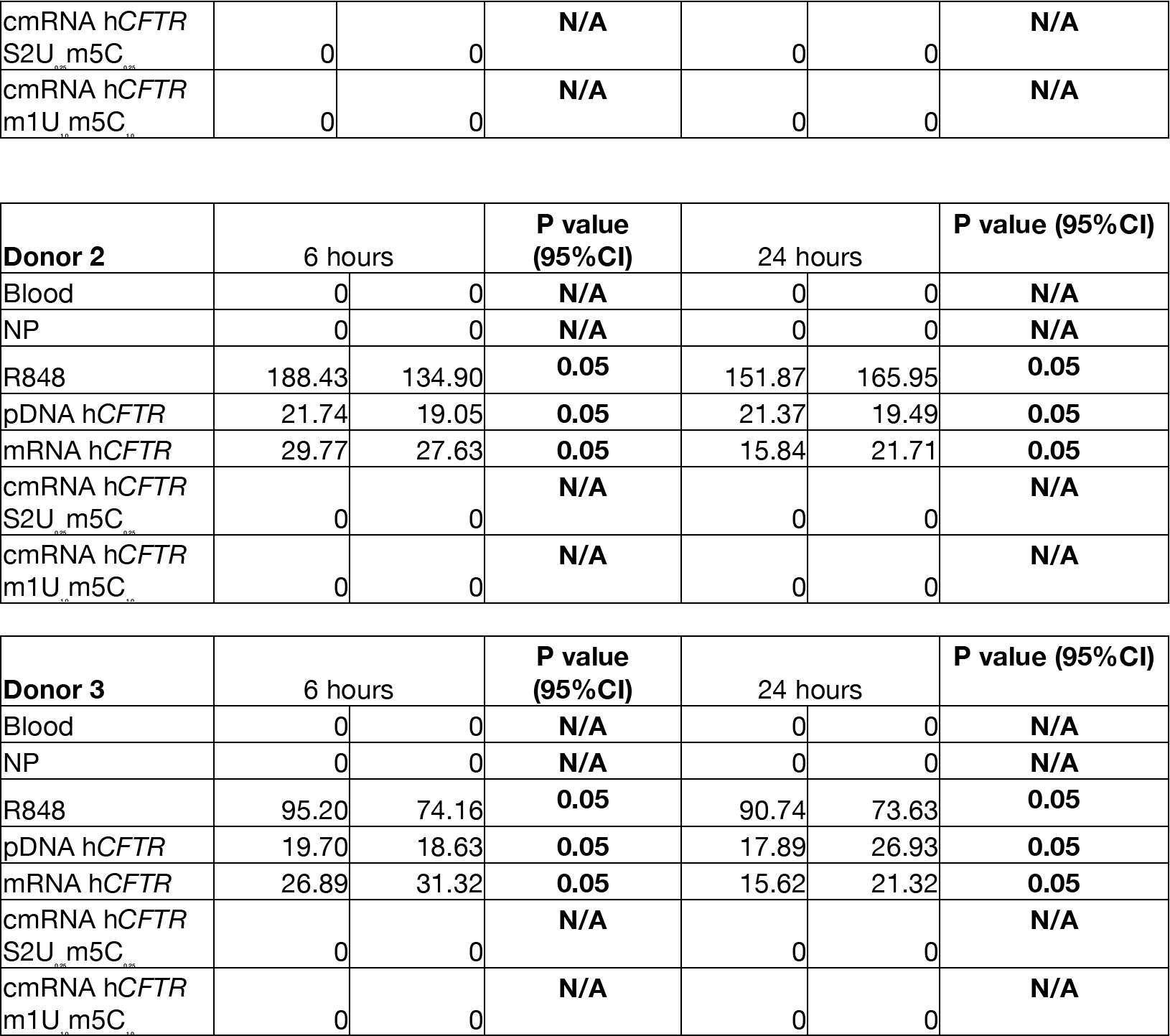

**Fig. 2D:**
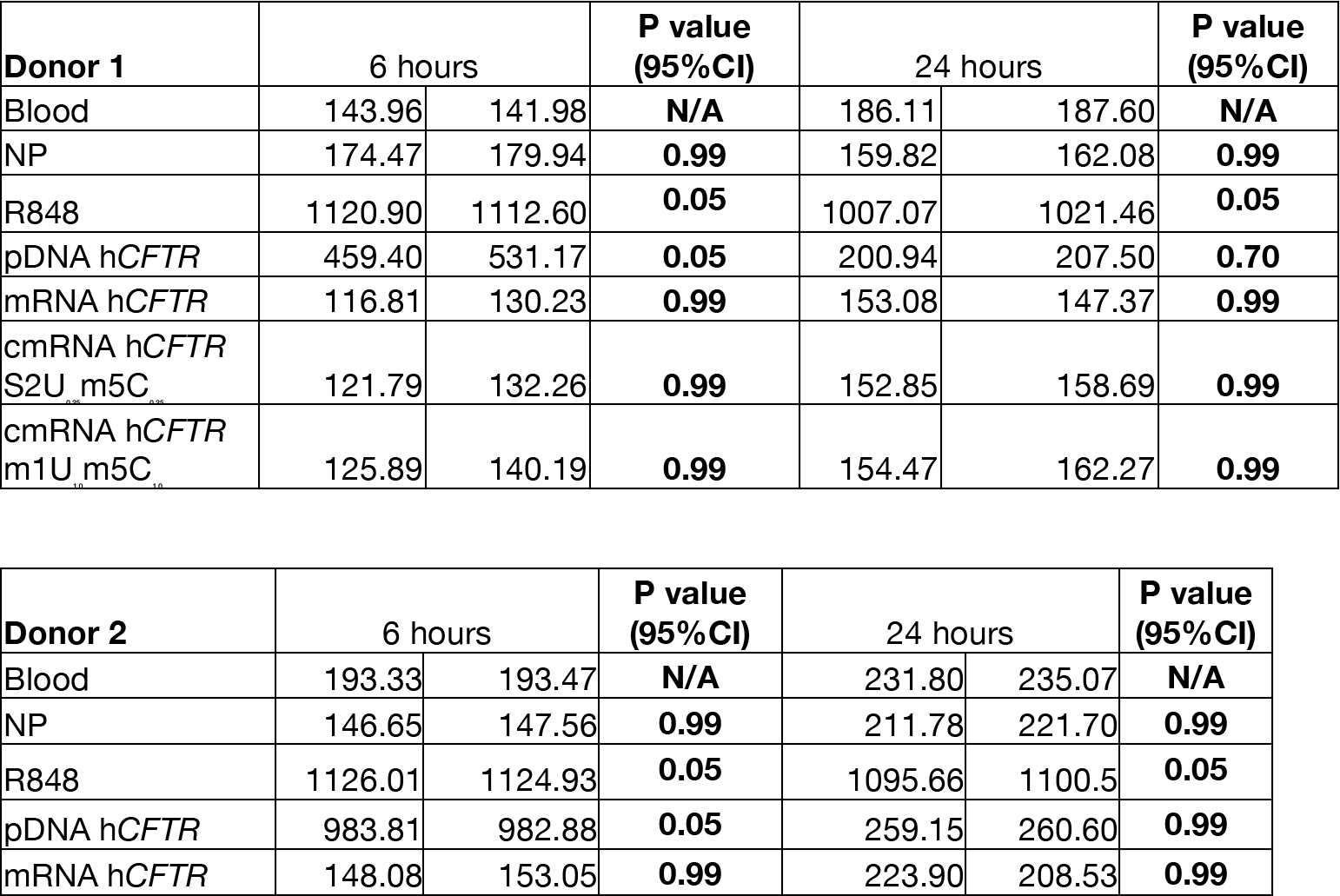
Whole blood assay INF-Alpha (*ex vivo*) **Donor (n=3) Source 14** **Statistic:** Wilcoxon-Mann-Whitney test and *P* ≤ 0.05(two-sided) was considered statistically significant

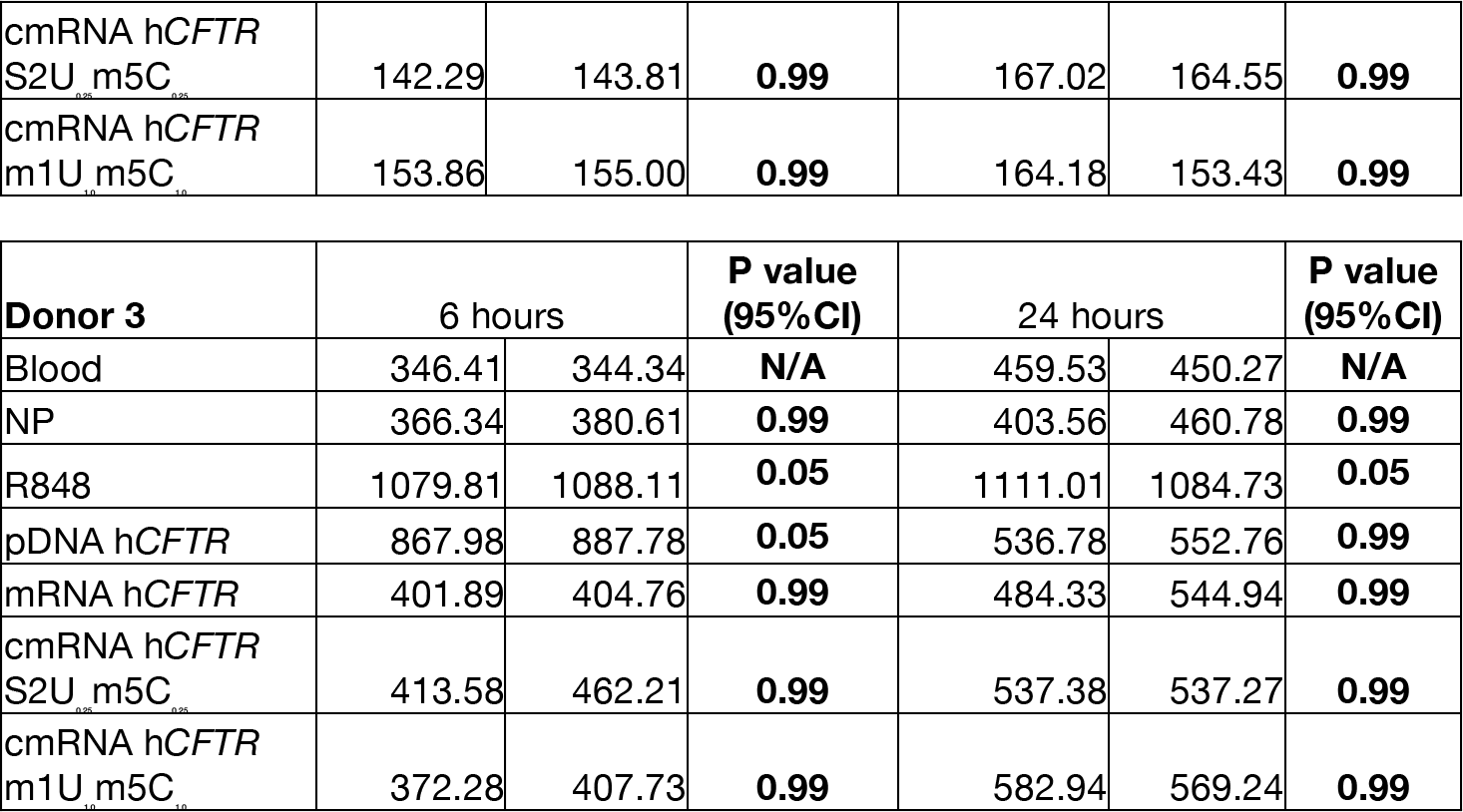

**Fig. 3B:**
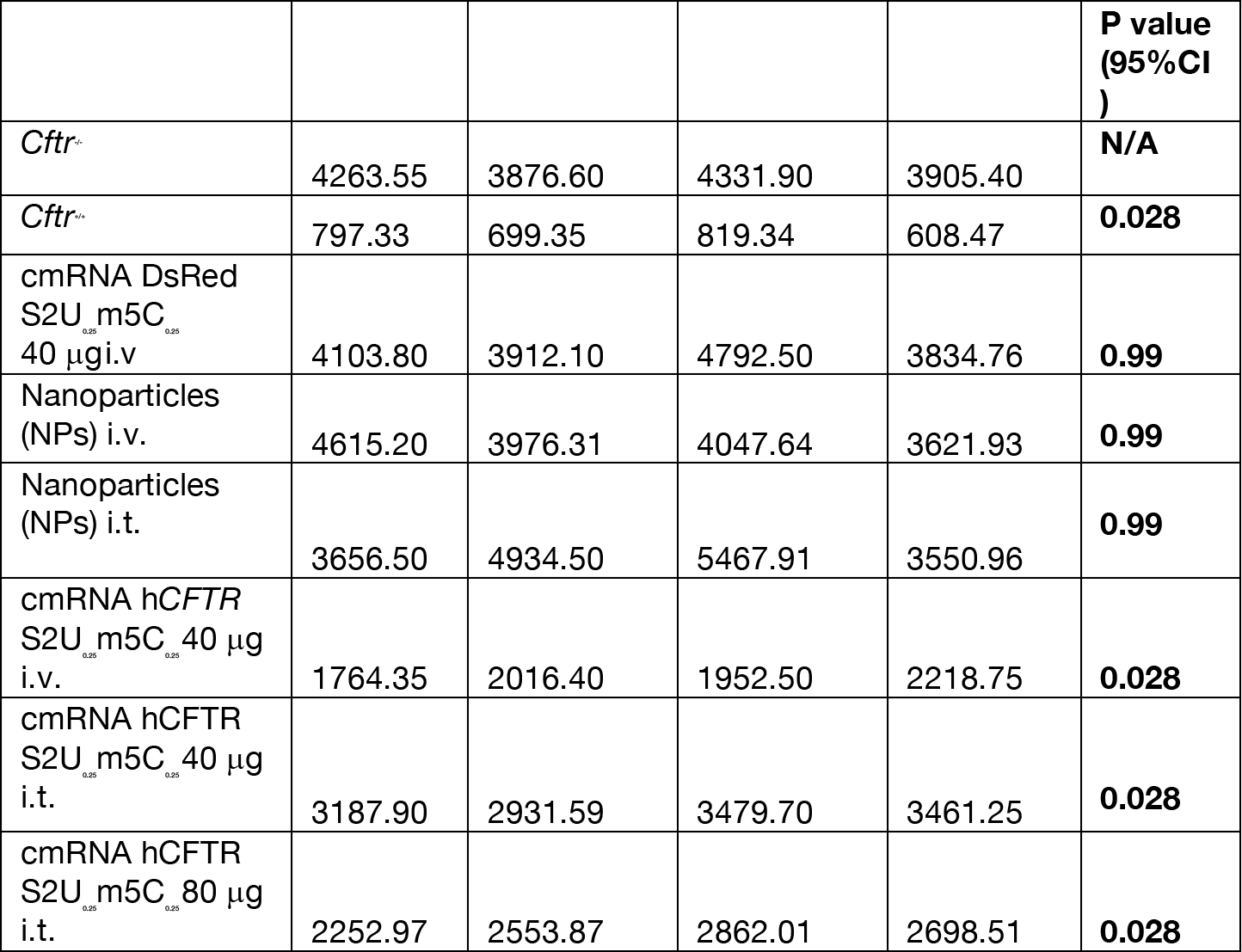
Salivary Chloride assay, **n=4: Source 15** **Statistic:** Wilcoxon-Mann-Whitney test and *P* ≤ 0.05(two-sided) was considered statistically significant

**Fig. 3C:**
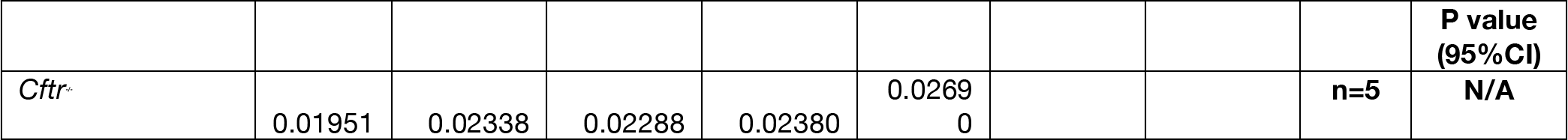
Lung Compliance. **Source Data 16** **Statistic:** Wilcoxon-Mann-Whitney test and *P* ≤ 0.05(two-sided) was considered statistically significant

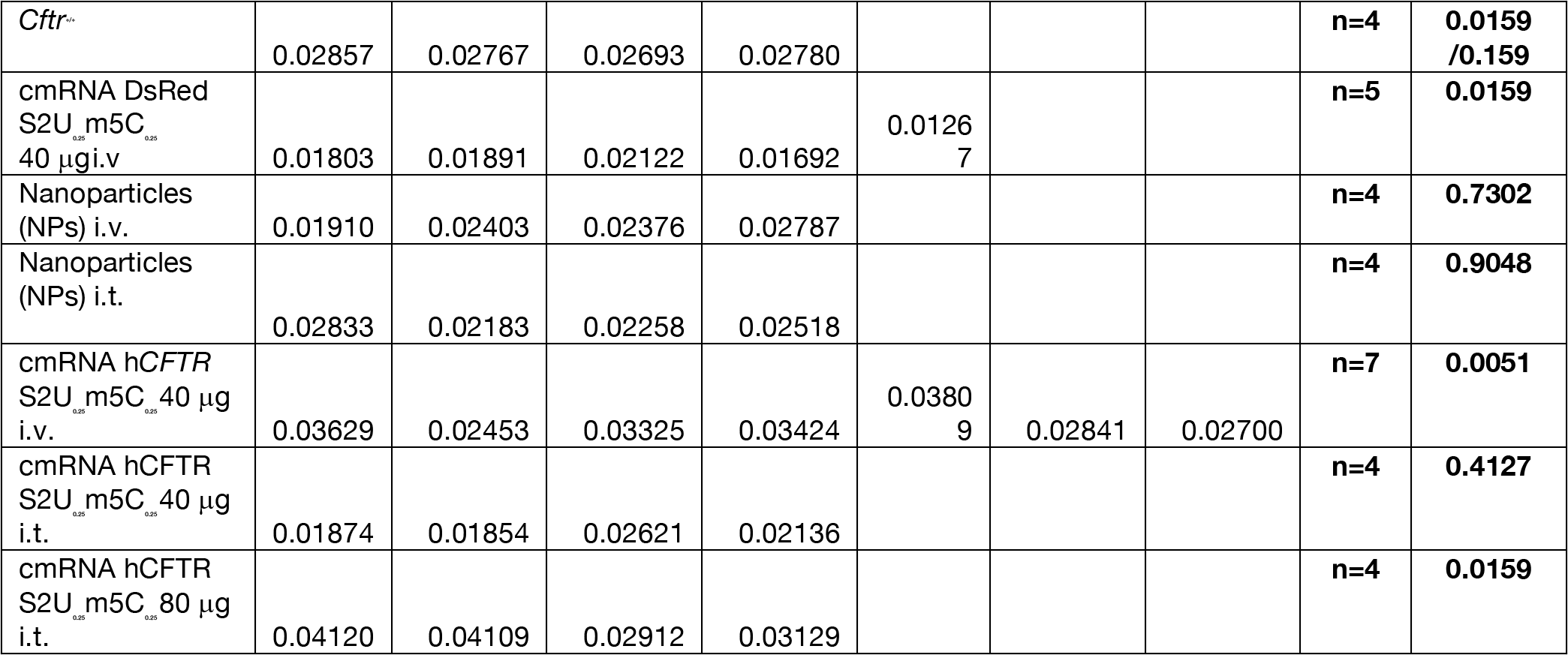

**Fig. 3D:**
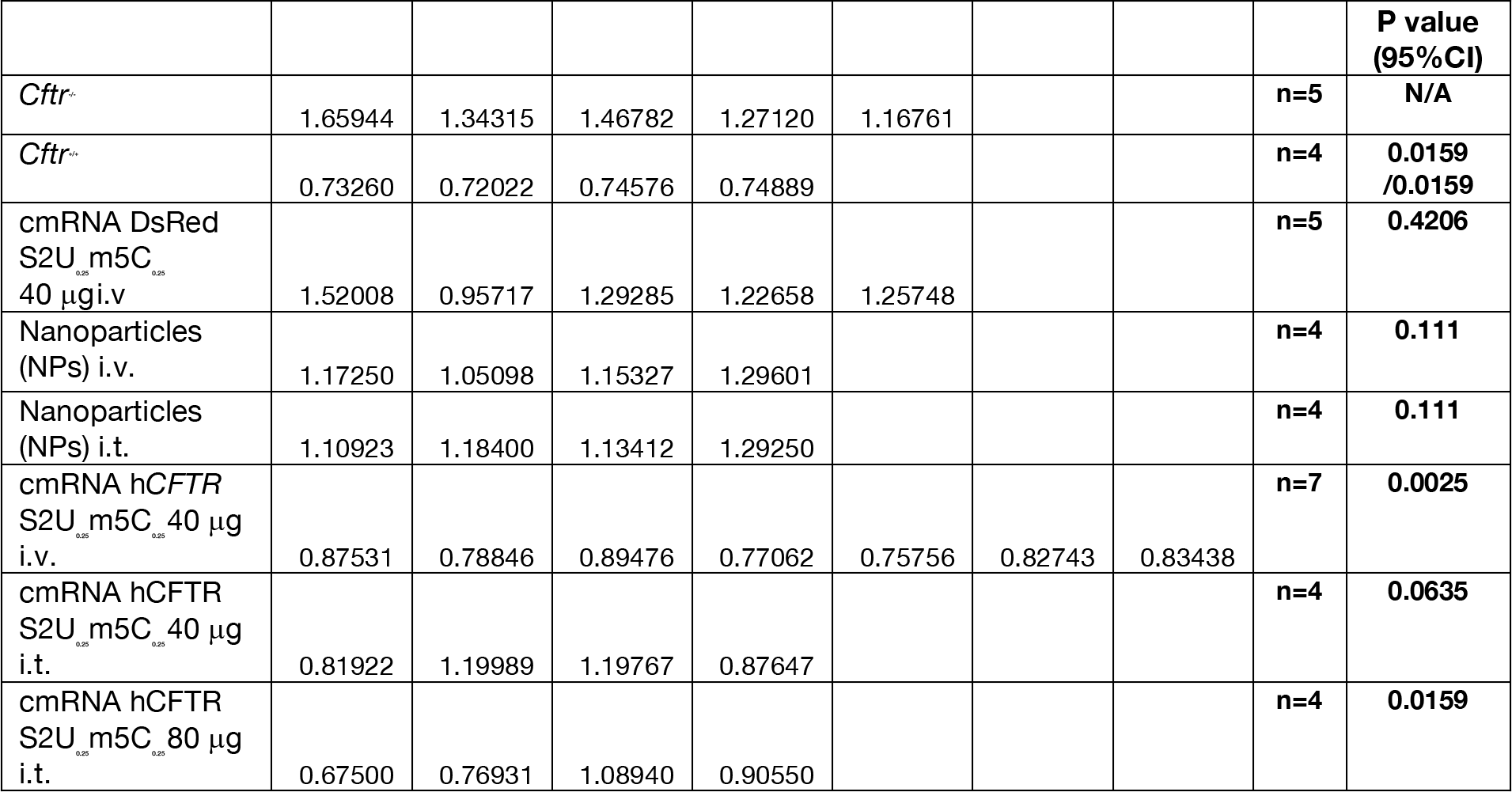
Lung Resistance. **Source 17** **Statistic:** Wilcoxon-Mann-Whitney test and *P* ≤ 0.05(two-sided) was considered statistically significant

**Fig. 3E:**
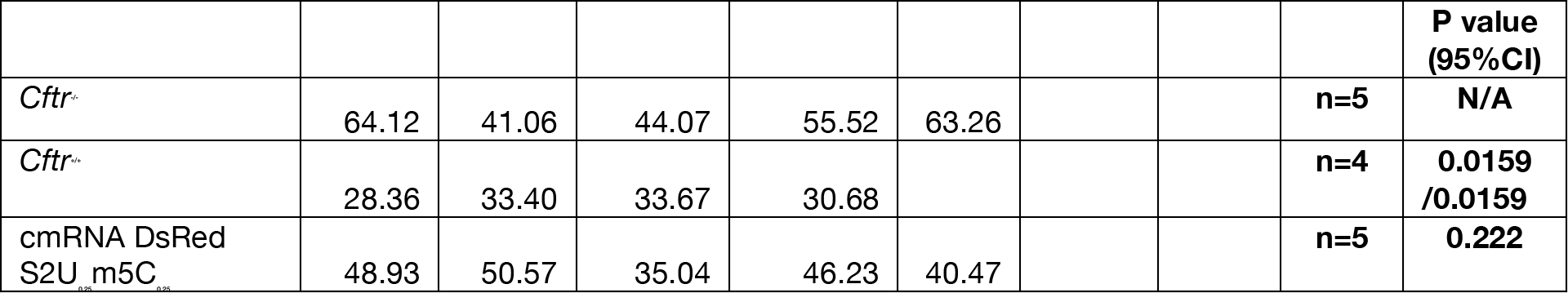
Tissue Elastance. **Source Data 18** **Statistic:** Wilcoxon-Mann-Whitney test and *P* ≤ 0.05(two-sided) was considered statistically significant

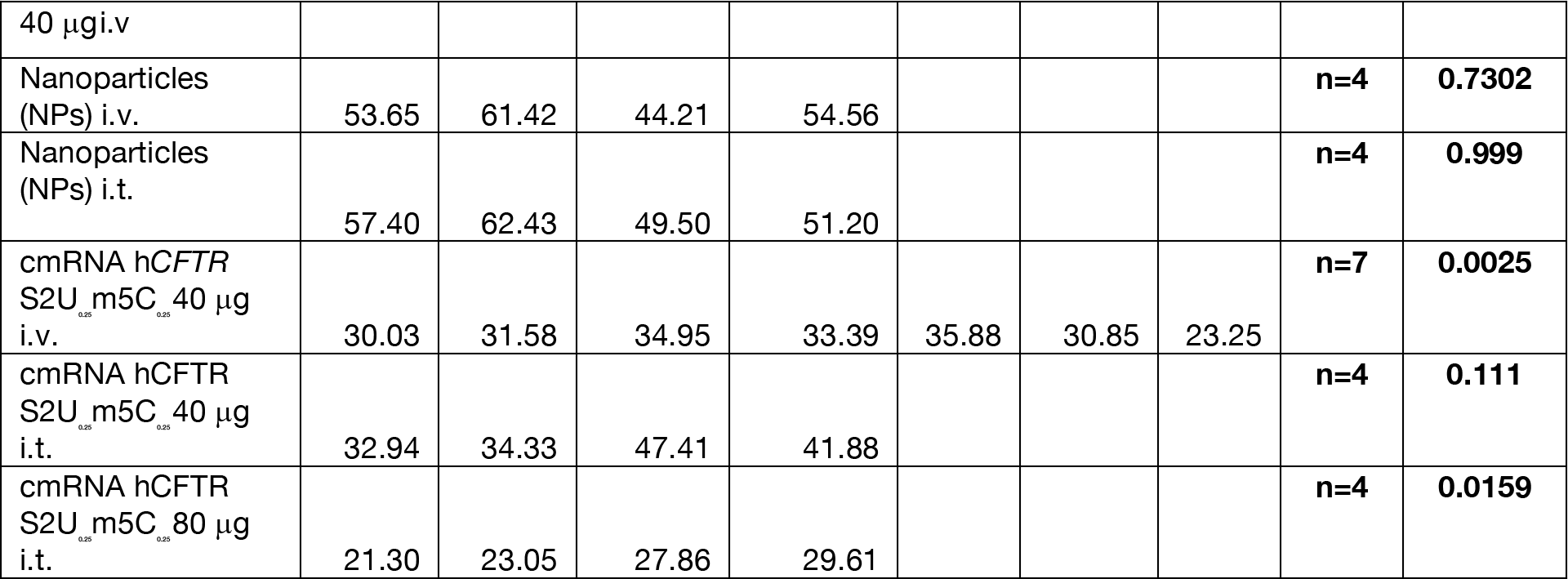

**Fig. 3F:**
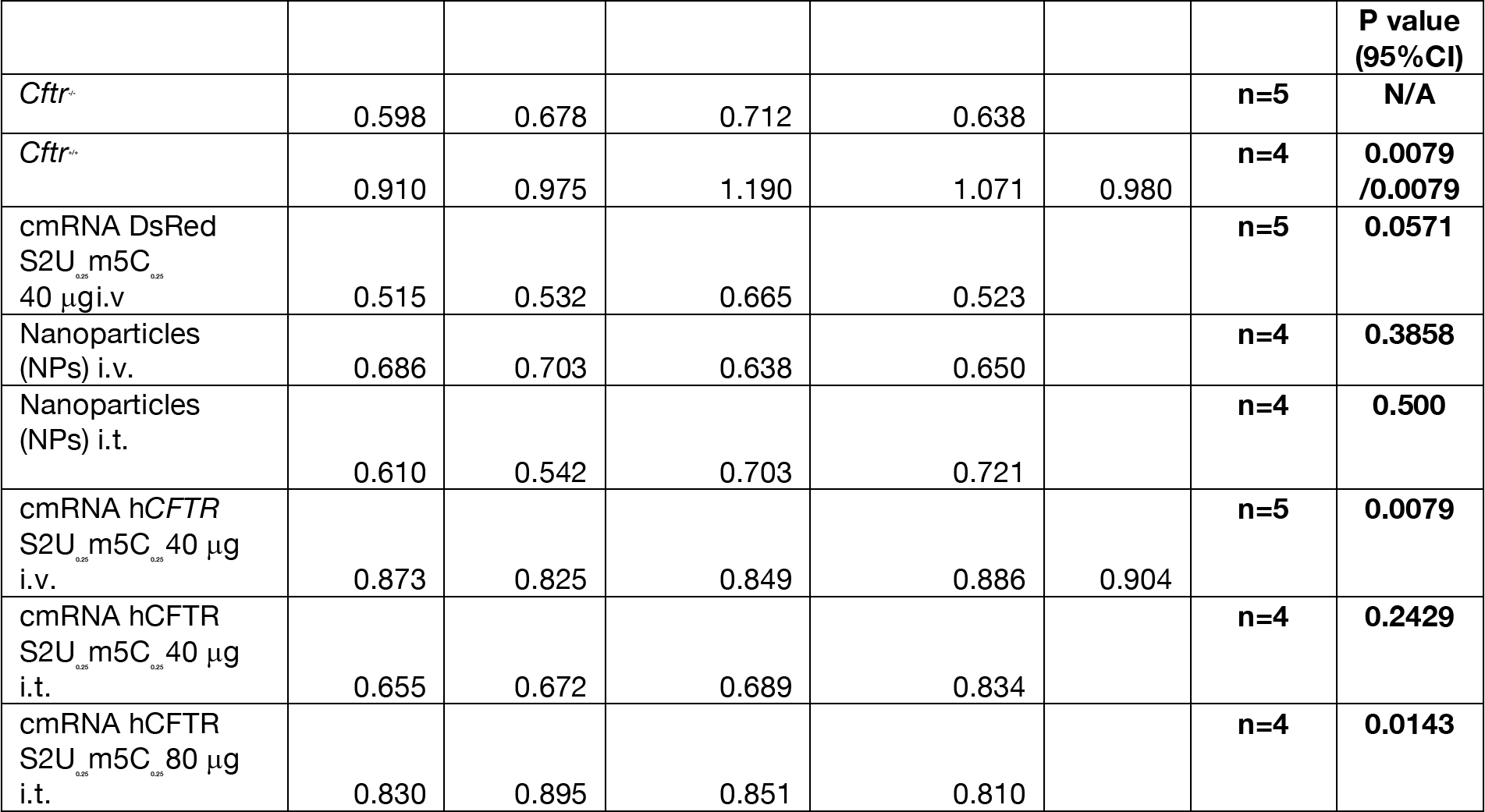
FEV. **Source Data 19** **Statistic:** Wilcoxon-Mann-Whitney test and *P* ≤ 0.05(two-sided) was considered statistically significant

**Fig. 4B:**
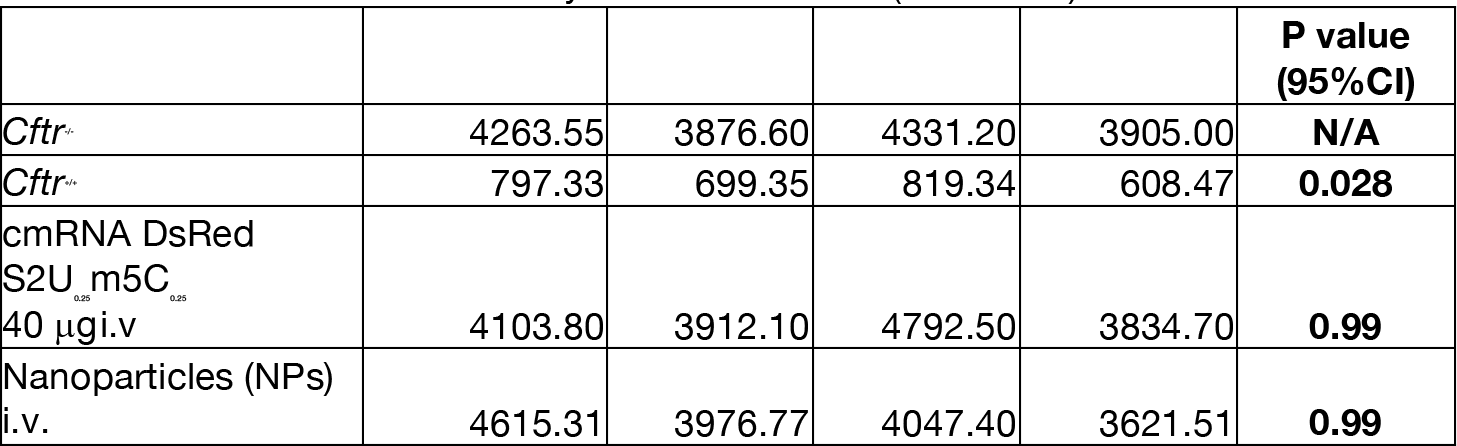
Salivary Assay, **n = 4: Source 20** **Statistic:** Wilcoxon-Mann-Whitney test and *P* ≤ 0.05(two-sided) was considered statistically significant

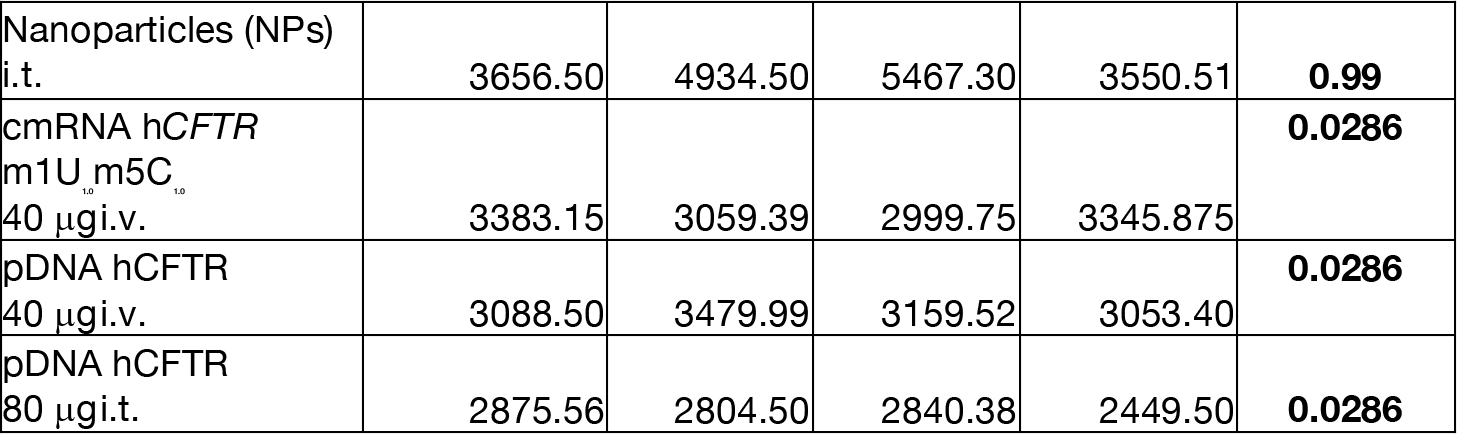

**Fig. 4C:**
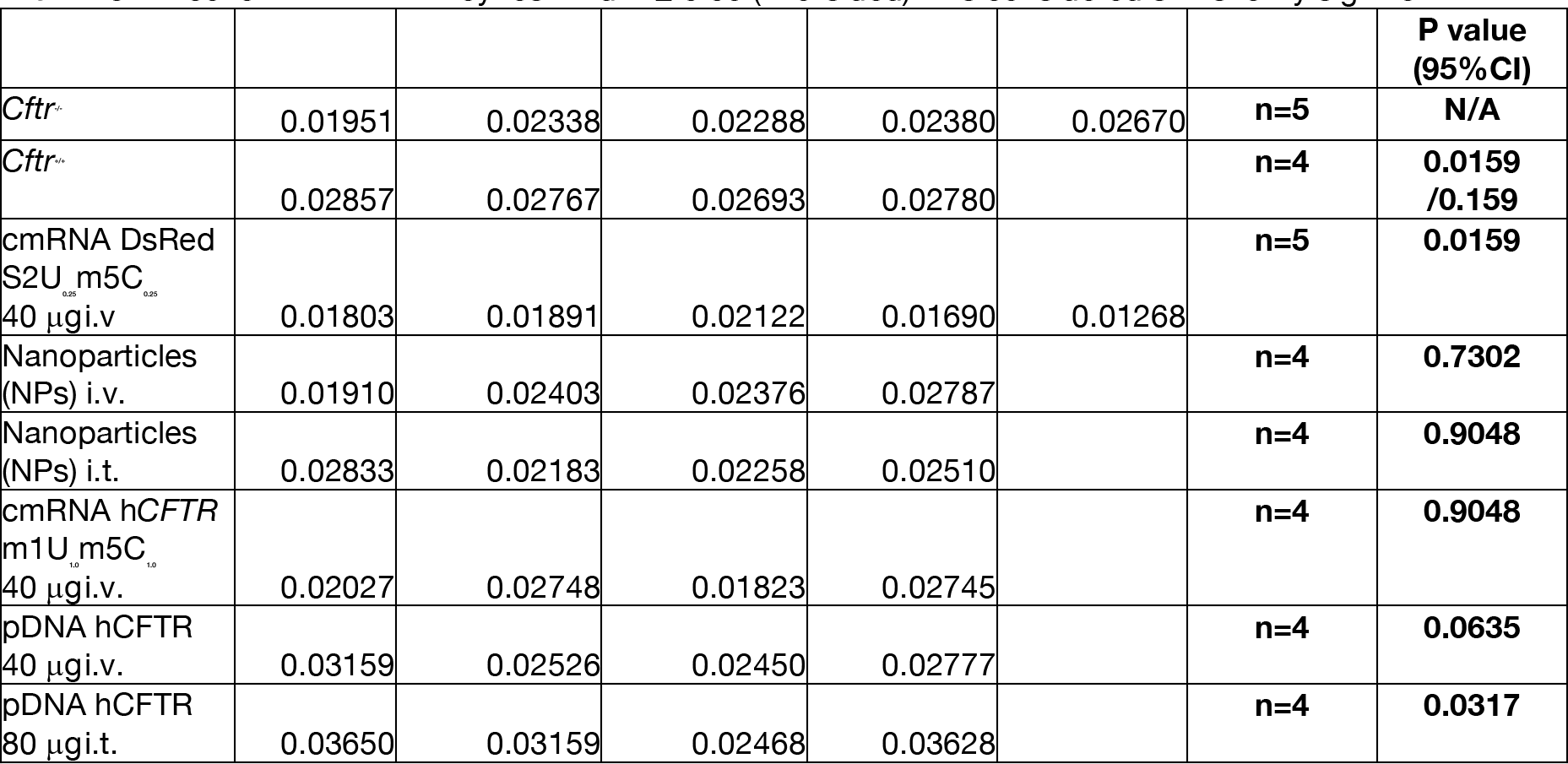
Lung Compliance. **Source Data 21** **Statistic:** Wilcoxon-Mann-Whitney test and *P* ≤ 0.05(two-sided) was considered statistically significant

**Fig. 4D:**
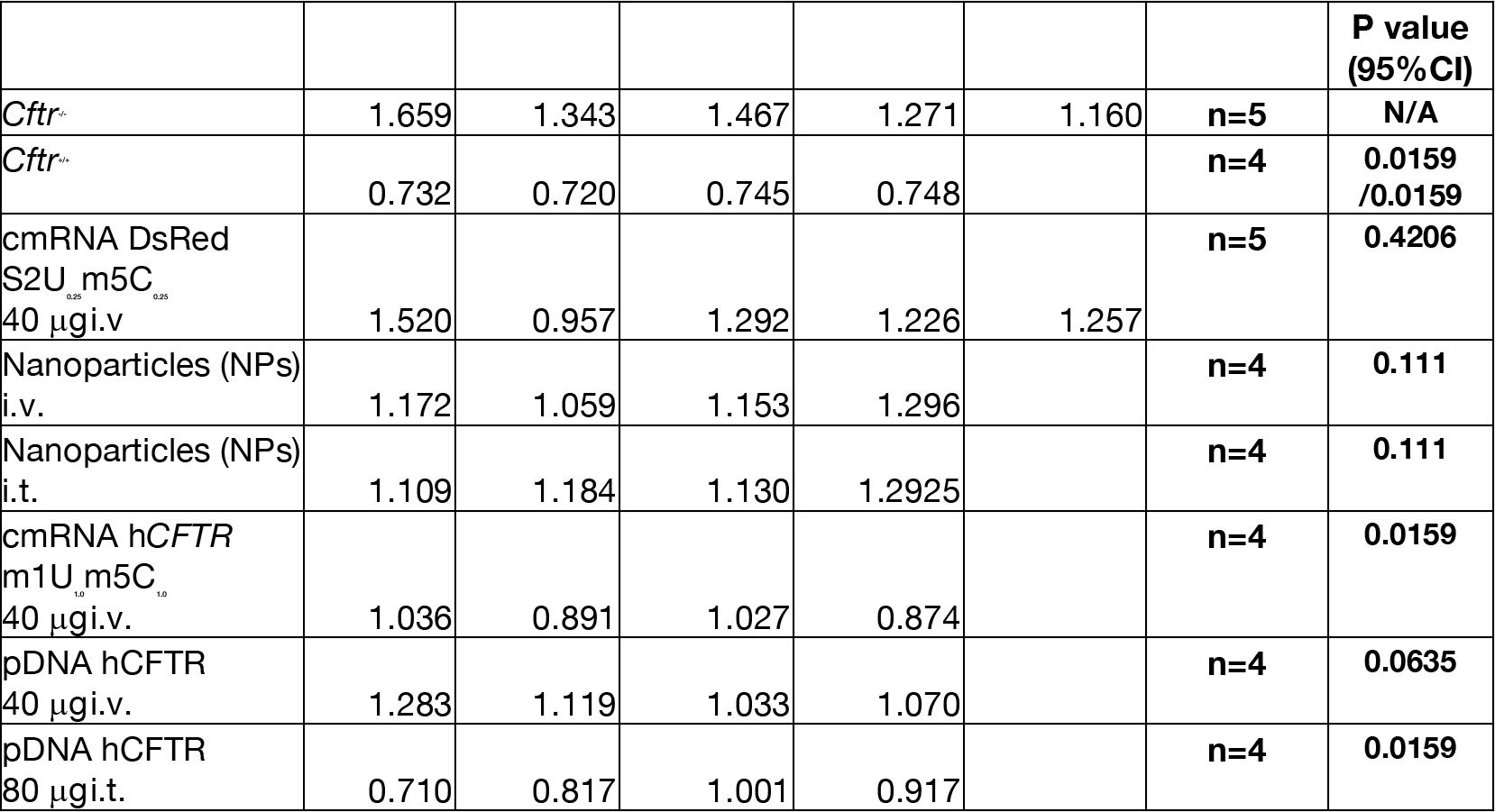
Lung Resistance. Source Data 21. **Statistic:** Wilcoxon-Mann-Whitney test and *P* ≤ 0.05(two-sided) was considered statistically significant

**Fig. 4E:**
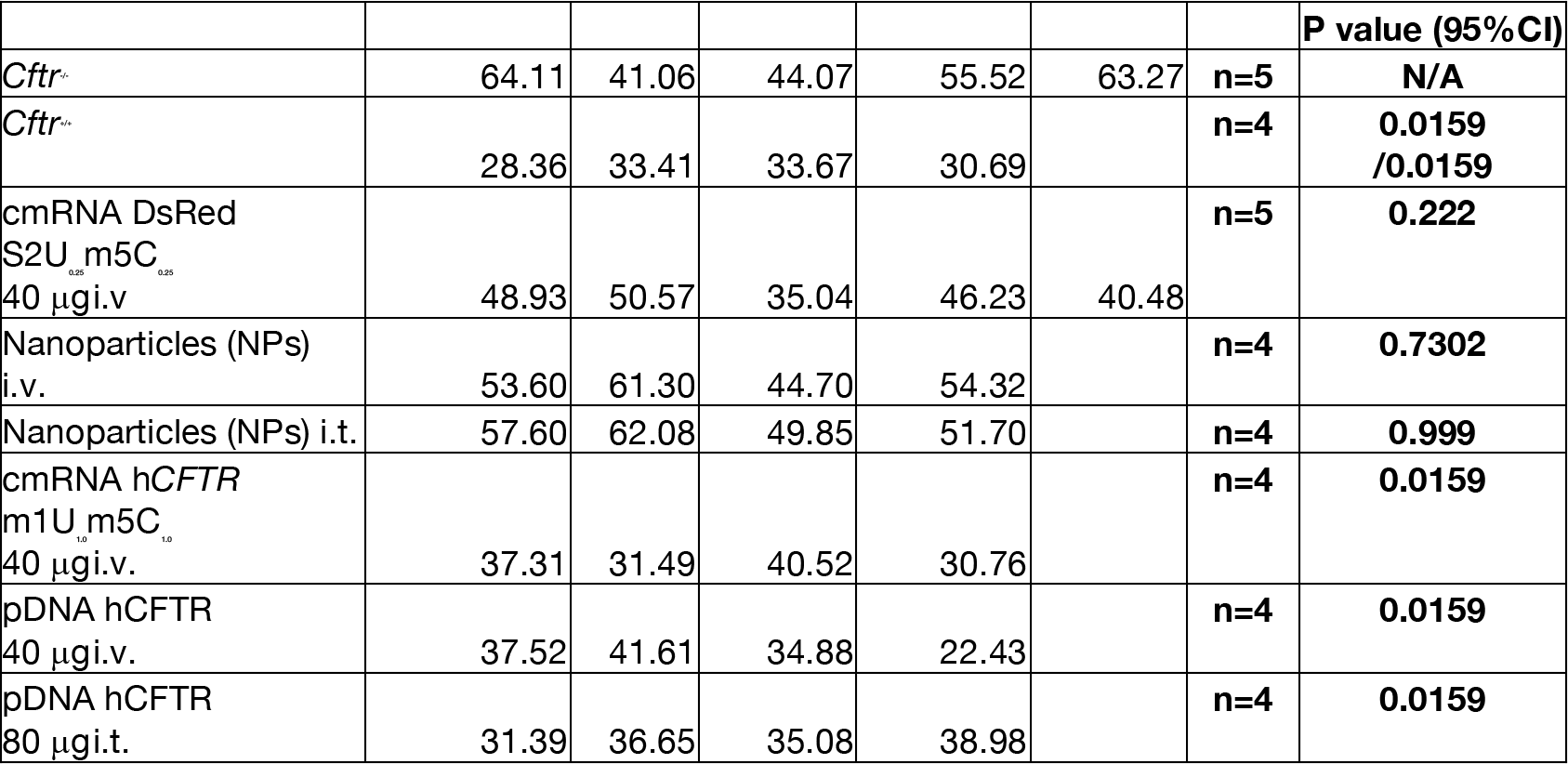
Tissue Elastance Source 22. **Statistic:** Wilcoxon-Mann-Whitney test and *P* ≤ 0.05(two-sided) was considered statistically significant

**Fig. 4E:**
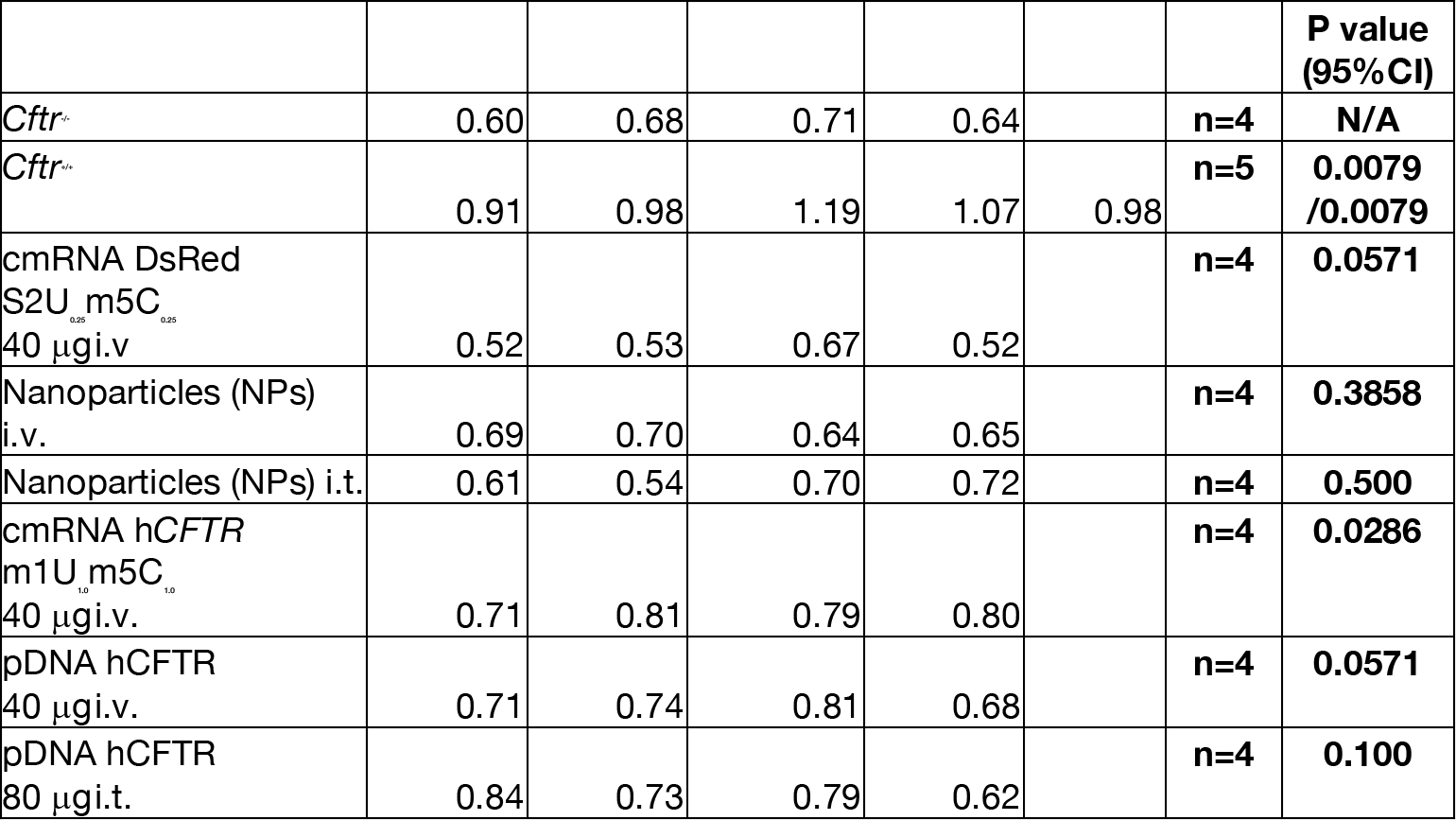
FEV_0.1_ Source 22 **Statistic:** Wilcoxon-Mann-Whitney test and *P* ≤ 0.05(two-sided) was considered statistically significant

